# The Novel Role of Midbody-Associated mRNAs in Regulating Abscission

**DOI:** 10.1101/2022.10.27.514111

**Authors:** Trey Farmer, Katherine F. Vaeth, Ke-Jun Han, Raeann Goering, J. Matthew Taliaferro, Rytis Prekeris

## Abstract

Midbodies (MBs) have been shown to function during telophase as a recruiting hub, especially for ESCRT-III complex subunits, to regulate the abscission step of cytokinesis. However, the molecular machinery governing specific protein targeting and activation at the MB remains poorly understood. Until recently, it was thought that abscission regulating proteins, such as ESCRT-III complex subunits, accumulate at the MB by directly or indirectly binding to the MB resident protein, CEP55. However, recent studies have shown that depletion of CEP55 does not fully block ESCRT-III targeting to the MB, and cells in CEP55 knock-out mice divide normally. Additionally, since MBs are microtubule-rich, proteinaceous structures, it is conceptually hard to imagine how large protein complexes, such as the ESCRT-III complex, can successfully diffuse into the MB from the cytosol in a rapid and highly regulated manner. Here, we show that MBs contain mRNAs and that these MB-associated mRNAs can be locally translated, resulting in the accumulation of abscission-regulating proteins. We also demonstrate that localized MB-associated translation of CHMP4B is required for its targeting to the abscission site and that 3′ UTR-dependent CHMP4B mRNA targeting to the MB is required for successful completion of cytokinesis. Finally, we identify regulatory *cis*-elements within RNAs that are necessary and sufficient for mRNA trafficking to the MB. Based on all this data, we propose a novel method of regulating cytokinesis and abscission by MB-associated targeting and localized translation of selective mRNAs.

## Introduction

The cell cycle is a key event during the development, growth, and reproduction of all organisms. Cytokinesis is the final stage of the cell cycle that results in the physical separation of two daughter cells. The mother cell divides by the formation of the cleavage furrow, leaving two daughter cells connected by a microtubule-rich intercellular bridge. Additionally, during ingression of the cleavage furrow, the central spindle microtubules are compacted to form a structure known as the midbody (MB). To complete cytokinesis, the intercellular bridge must be cleaved on one or both sides of the MB, via the process known as the abscission (Figure 1A).Abscission is a highly regulated event, and it is now well established that the MB plays a key role in regulating completion of cytokinesis by recruiting and activating abscission-mediating proteins, such as the Endosomal Sorting Complex Required for Transport (ESCRT-III) complex, as well as several regulators of the abscission checkpoint^1, 2^. While targeting and activation of abscission regulators at the MB is required for completion of cytokinesis, the mechanisms governing this process remain to be fully defined.

**Figure 1.**
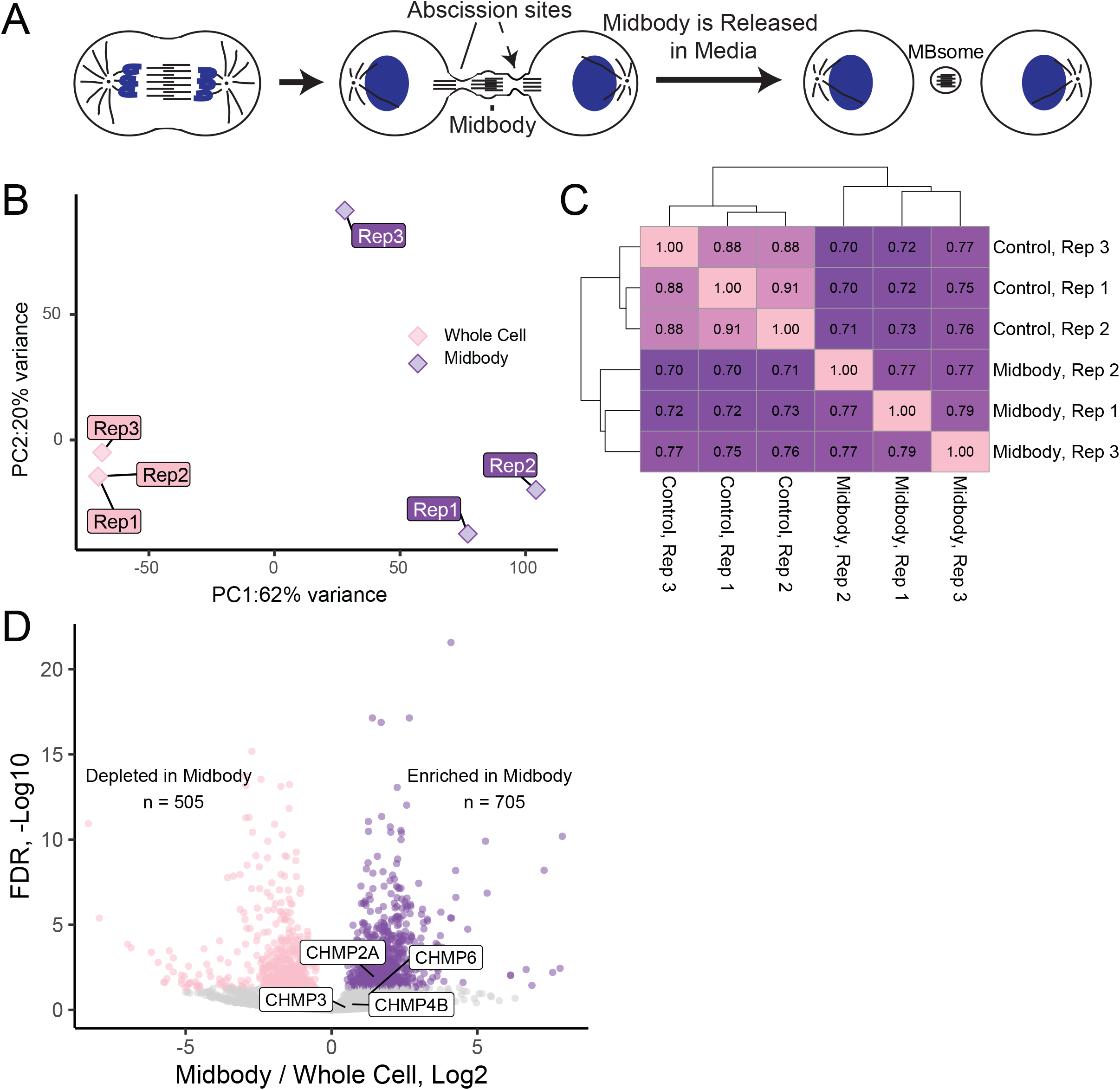
MBsomes contain a distinct subset of mRNAs. (A) Schematic representing MB formation and MBsome release into the media. MBsomes from the media were isolated for RNAseq analysis presented in this study. (B) Principal component analysis representing the difference between the 3 control and the 3 MB RNAseq datasets. (C) Correlation analysis representing the correlation between the various control and MB RNAseq datasets. (D) Volcano plot showing the significantly depleted (pink) and significantly enriched (purple) mRNAs found within MBsomes from 3 RNAseq datasets.

Previous studies have shown that several members of the ESCRT-III machinery are recruited to the MB just before initiation of the abscission^3–6^. Specifically, it was shown that TSG101 recruits Charged Multivesicular Body Proteins (CHMPs, components of the ESCRT-III complex)^7^ to the MB during telophase. In addition to TSG101, ALIX (also known as PDCD6IP) also contributes to the direct recruitment of the ESCRT-III complex to the MB^1, 4, 8^. Upon recruitment to the MB, the ESCRT-III components oligomerize (CHMP4B and CHMP2A or CHMP3 and CHMP6) into spiral structures with progressively smaller diameters towards the abscission site^9–11^, the process that is thought to be responsible for the eventual abscission of the intercellular bridge to complete cytokinesis and form two daughter cells^6, 9, 10^. Inhibiting MB recruitment of ESCRT-III results in abscission defects, demonstrating the importance of appropriately regulating ESCRT-III protein accumulation and activation at the MB^5, 6, 12–15^.

Until recently, it has been thought that TSG101 accumulates at the MB by binding to the MB resident protein, CEP55^3, 4, 6, 9, 10, 16, 17^. This binding was presumed to tether the TSG101 at the MB, gradually leading to ESCRT-III accumulation before initiation of abscission. However, recent studies have shown that depletion of CEP55 does not fully block ESCRT-III targeting to the MB, and CEP55 knock-out mice are able to grow and develop^18^. Finally, CEP55 is a protein present only in vertebrates, thus, while CEP55 clearly contributes to ESCRT-III recruitment to the MB in vertebrates, it does not appear to be absolutely required for abscission. Additionally, the MB and intercellular bridge are microtubule-rich, diffusion-limited structures, raising a question of how large protein complexes, such as the ESCRT-III complex, can rapidly diffuse via the intercellular bridge into the MB from the cytosol in a timely and efficient manner.

Recent work suggested that in addition to regulating abscission, secreted post-mitotic MBs (also known as MBsomes, MB remnants, or Flemmingsomes) can be internalized by other cells where they regulate cell proliferation and differentiation^8, 19–24^. It was proposed that these MBsomes may mediate cell-to-cell transfer of various signaling molecules in a manner similar to exosomes and extracellular vesicles^23, 24^. Consequently, several laboratories, including ours, have analyzed the proteome of secreted MBsomes. Interestingly, these studies have shown that, in addition to various signaling proteins, MBs are also enriched in ribosomal and translation initiation factor proteins^8, 23–25^, raising an intriguing possibility that MB-associated mRNA targeting and localized translation may mediate various mitotic and post-mitotic MB functions.

In this study we show, for the first time, that a specific subset of mRNAs is indeed targeted to and enriched at the MBs. We also demonstrate that MBs contain active ribosomes and that MB-associated protein translation is needed for successful completion of the abscission. Importantly, by using a combination of RNAseq, qPCR, and smFISH, we demonstrate that mRNA encoding ESCRT-III components, specifically CHMP4B, CHMP2A, CHMP3, and CHMP6, are all enriched at the MB, and that MB-associated translation is needed for CHMP4B accumulation at the MB during telophase. Additionally, characterization of CHMP4B mRNA revealed that the 3′ UTR is critical for efficient CHMP4B mRNA targeting to the MB. Finally, we identify GA-rich localization elements within 3′ UTR that are necessary and sufficient to target mRNA to the MB. Collectively, this study proposes a novel mechanism of regulating cytokinesis and abscission by MB-associated targeting and localized translation of selective mRNAs that include, but are not limited to, the members of ESCRT-III complex. These findings lay a foundation for future studies in determining the molecular mechanisms regulating mRNA targeting and translation at the MB, and future analyses of the roles that this mRNA targeting plays in regulating abscission and post-mitotic MBsome functions.

## Results

### Identification of Midbody-Associated mRNAs

Recent studies have shown that MBs are enriched in RNA-binding proteins^8, 23–25^, leading to the intriguing idea that MBs may contain mRNAs, and that local translation at the MB may regulate various MB functions. To test this hypothesis, we first set out to determine whether specific mRNAs actually accumulate at the MBs and began by isolating MBsomes from tissue culture media as previously described^26^ (Figure 1A). We then prepared oligo-dT-based mRNA libraries from MBsomes and matched whole cell RNA samples, analyzing them using RNAseq (Supplemental Table 1). To assess the reproducibility of the RNA content of our MBsome preparations and their relationship to whole cell RNA samples, we used principal components analysis and hierarchical clustering (Figure 1B, C). With both approaches, we found that the MB RNA samples were highly similar to each other, indicating the reproducible nature of their RNA contents. Importantly, they were well separated from the whole cell RNA samples, suggesting that the MB RNA samples contained an mRNA composition distinct from that found in the whole cell samples. Overall, the RNAseq analysis identified 16,922 mRNAs that can be detected in the MBsome. From those, 505 mRNAs were significantly depleted and 705 mRNAs were significantly enriched in the MBsomes as compared to the whole cell RNA samples (Figure 1D; Supplemental table 1).

MBsomes are produced exclusively by mitotic cells. Our observed MBsome-enriched RNAs, then, could therefore simply reflect the RNA content of cells that produced MBsomes (i.e. cells undergoing mitosis) rather than RNAs specifically trafficked to the midbody. To assess whether our observed MBsome RNA enrichments reflected MBsome-localized RNAs or the general RNA content of mitotic cells, we compared our data to RNAseq data from unsynchronized HeLa cells sorted for cell cycle phase using Fucci reporter expression^43^. For all RNAs, we calculated their relative enrichment in cells in G2/M phase compared to cells in G1. We found that our MBsome-enriched RNAs were significantly less abundant in G2/M cells compared to G1 cells (Figure S1A). Similarly, MBsome-enriched RNAs were less abundant in cells in S phase compared to G1 cells (Figure S1B). From these data, we conclude that RNA enrichments in MBsomes are not due to their increased abundance in mitotic cells.

Given the MB’s connection to the mitotic spindle, we also wondered whether MBsome-enriched RNAs were generally localized to the mitotic spindle or were specific to the MB. To test this, we analyzed RNAseq data in which spindle-associated RNAs were identified through differential centrifugation^44^. We found that MBsome-enriched RNAs were not spindle-enriched, indicating that these two RNA populations are likely distinct (Figure S1C).

To determine the possible function of these MB-enriched mRNAs, we performed a Gene Ontology (GO) Enrichment Analysis (Figure 2A) and found that mRNAs encoding for proteins that regulate cell cycle and microtubule dynamics are enriched at the MBsomes (Figure 2A). We also determined that mRNAs of several abscission-regulators, such as Citron Kinase, CDK11A, CDK11B, and AKTIP, are also enriched in the MBsomes (Figure 2B). Importantly, we found that mRNAs of the ESCRT-III complex proteins, CHMP4B, 2A, 3, and 6, were all enriched at the MBsome (Figure 2B-D). This is especially noteworthy as CHMP4B/2A and CHMP3/6 are known to co-polymerize and drive the final abscission step^9–11^ (Figure 2D), suggesting that localized translation of ESCRT-III proteins may drive their accumulation at the MB and may also be required for the abscission step of cytokinesis.

**Figure 2.**
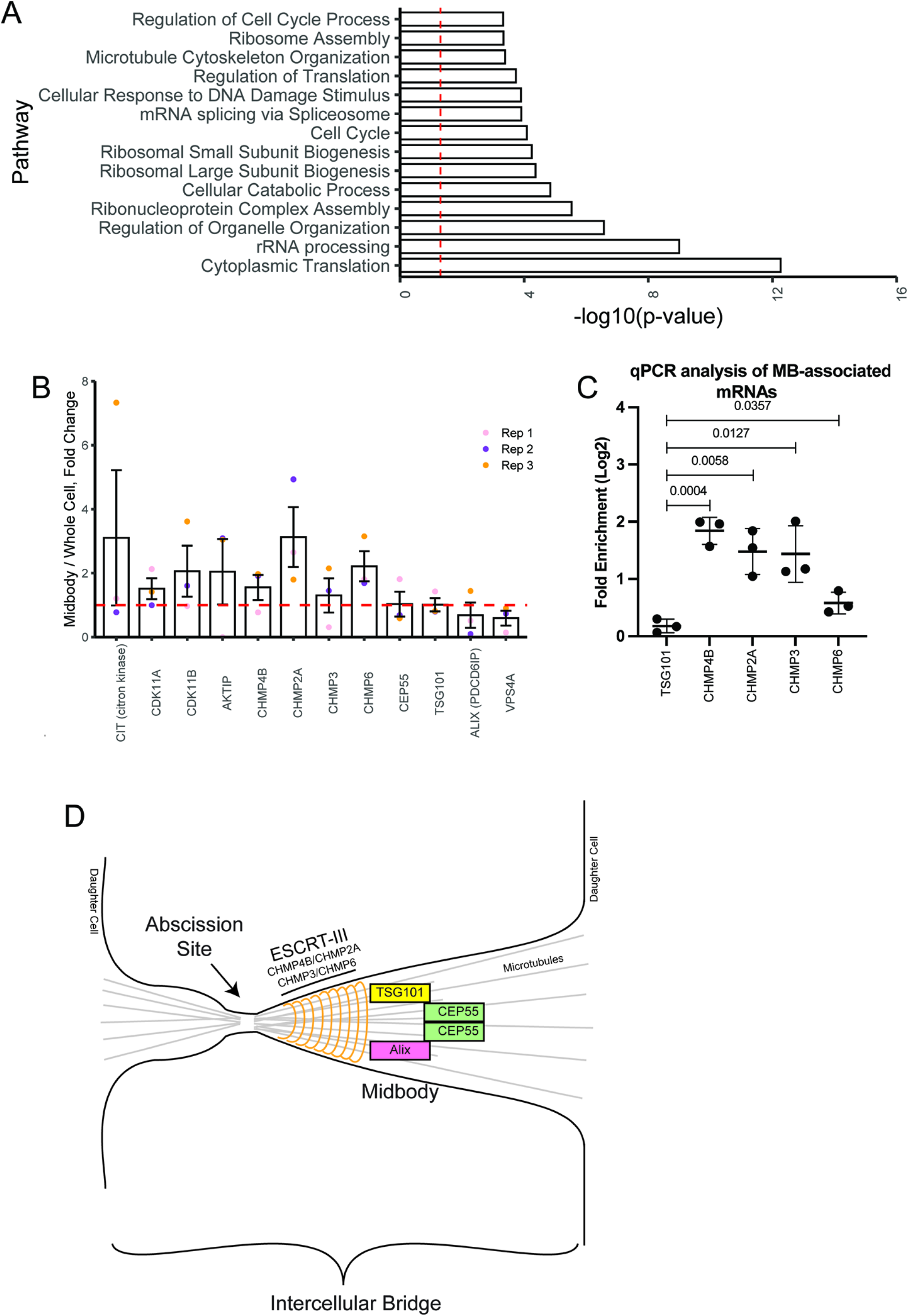
**MBsomes are enriched in mRNAs encoding proteins that regulate translation and cell cycle.** (A) The Gene Ontology (GO) pathway analysis tool was used to determine the pathways associated with the enriched MB-associated mRNAs. (B) MB enrichment of mRNAs encoding known abscission regulators. Data shown presents means and standard deviations derived from three separate RNAseq analyses. Dashed line marks the mRNA levels present in the whole cell transcriptome. (C) Whole cell or MBsome-associated mRNAs were isolated from GFP-MKLP1-expressing HeLa cell line. The cDNA was subject to RT-qPCR using TSG101, CHMP4B, CHMP2A, CHMP3, CHMP6, or GAPDH (control) specific primers and represented as normalized values. Statistical analysis is represented with a p-value. (D) Schematic representing key regulators of abscission and how they relate to each other in the intercellular bridge.

Interestingly, not all mRNAs of genes associated with abscission were found to be enriched at the MBsomes. For example, CEP55, TSG101, ALIX, and VPS4 mRNAs, while present in MBsomes, were not enriched as compared to the whole cell transcriptome (Figure 2B-C). This suggests that only a subset of abscission regulating proteins, specifically proteins that act during final steps of the abscission, may be controlled by local translation. Consistent with this hypothesis, CEP55, TCG101, and ALIX regulate early steps of the abscission, namely recruiting and tethering ESCRT-III to the MB (Figure 2D). In contrast, all ESCRT-III components are recruited during late telophase, just before completion of the abscission, suggesting an intriguing possibility that the MB targeting of proteins during telophase may depend on localized translation of specific mRNAs after MB formation. In the rest of the study, we will test this hypothesis by focusing on the machinery mediating MB targeting and translation of CHMP4B mRNA.

### Midbodies contain the molecular machinery required for protein translation

Our GO analysis (Figure 2A) demonstrated that numerous MB-associated mRNAs encode for proteins that regulate ribosomal assembly and function. This further supports the idea that MBs contain ribosomal machinery, which in turn can mediate local translation. Due to this finding, and the fact that CHMP4B mRNA is enriched in the MBs, we hypothesized that local translation of CHMP4B and other ESCRT-III subunits may contribute to ESCRT-III accumulation at the MB, therefore regulating the abscission. For this hypothesis to be correct, MBs must contain CHMP4B mRNA and translation associated ribosomal machinery, as well as be sites of active protein translation.

To test this idea, we first visualized CHMP4B mRNA using single molecule inexpensive RNA *in situ* hybridization (smiFISH)^27^. Consistent with our hypothesis that CHMP4B may be translated at the MB, we found that CHMP4B mRNA is present within the intracellular bridge during telophase (see arrows in Figure 3A) in over 60% of dividing HeLa cells (Supplemental Figure 2C-D). To validate the specificity of our CHMP4B smFISH probes, we depleted CHMP4B RNA using siRNA and observed a corresponding decrease in the number of observed CHMP4B RNA molecules per cell (Figure 3B).

**Figure 3.**
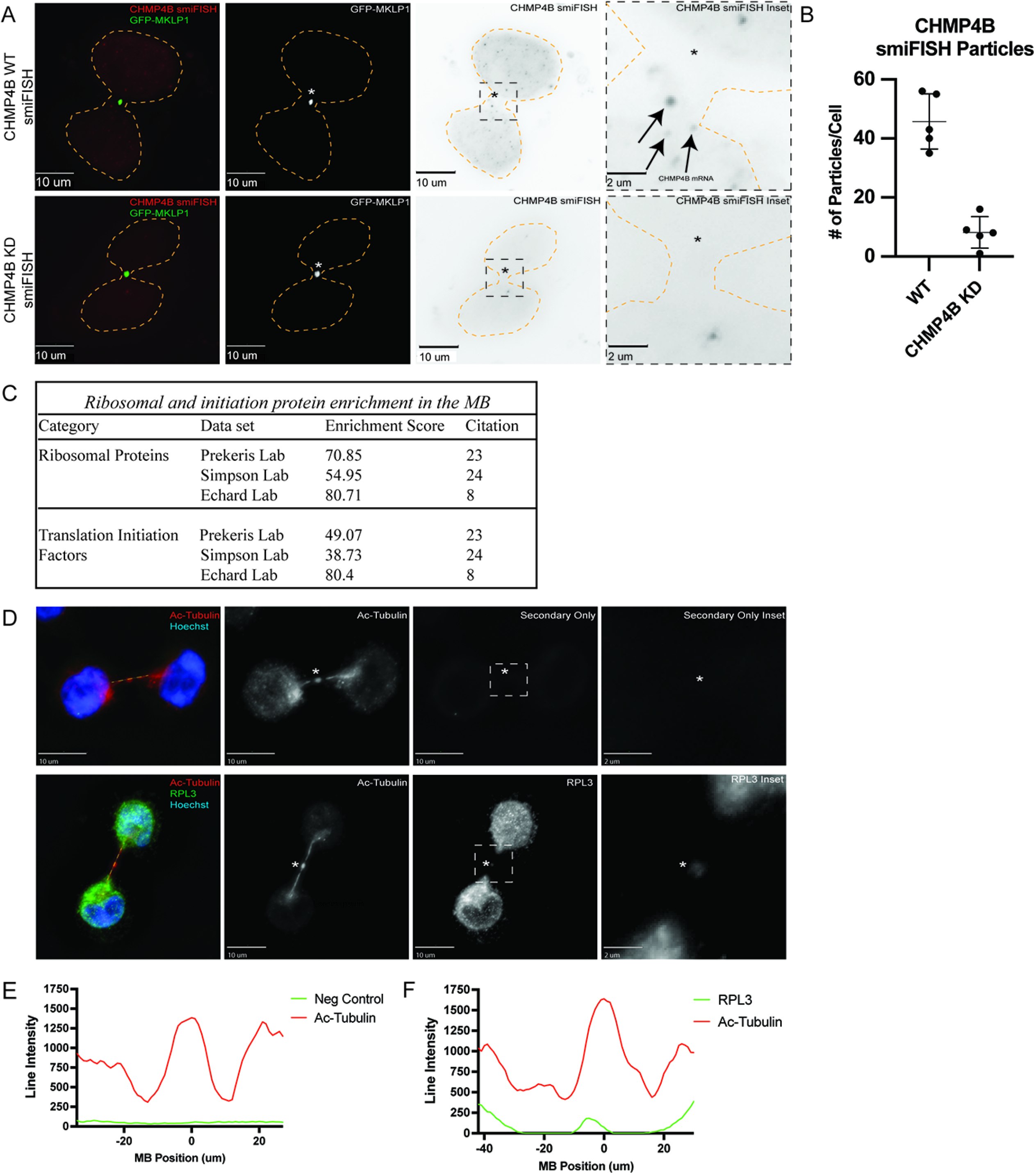
**Midbodies contain molecular machinery required for protein translation.** (A) GFP-MKLP1 expressing HeLa cells were either mock treated or treated with CHMP4B siRNA for 72 h and subjected to staining with smFISH probes against CHMP4B. Arrows in the inset points to CHMP4B mRNA particles. The cell outline is marked in yellow with the asterisks marking the MB. The black dashed square represents the region of the image used for the inset. (B) ImageJ was used to count the number of CHMP4B smFISH particles in the mock or CHMP4B siRNA treated cells. Each dot represents a single cell. (C) The Database for Annotation, Visualization, and Integrated Discovery (DAVID) was used to determine that ribosomal proteins and translation initiation factors are enriched in all 3 published MB proteomes. (D) HeLa cells were fixed and subjected to immunostaining with anti-acetylated-tubulin (red) and anti-RPL3 (green) antibodies. The line used for the line intensity graphs is marked in yellow and the asterisks mark the MB. The white dashed square represents the region of the image used for the inset. (E-F) Line intensity graphs representing the intensity of acetylated-tubulin (red) and RPL3 (green). Negative control was derived from samples that were treated with secondary antibodies but without anti-RPL3 antibodies.

There are multiple mechanisms that can drive mRNA accumulation at specific subcellular compartments. One is direct targeting of mRNA that is independent of translation, and another is co-translational targeting of mRNA that relies on the targeting of the nascent peptides (rather than mRNA) that emerge from the ribosome during translation. In this case, mRNA is indirectly accumulated to the specific subcellular compartment as the result of protein-dependent targeting of the entire mRNA-associated ribosome. To differentiate between these two possibilities, we briefly treated the cells with puromycin, a known inhibitor of protein translation that dissociates ribosomes from their mRNA substrates. As shown in Supplemental Figure 2A,C-D, puromycin treatment did not block CHMP4B mRNA accumulation at the MB, suggesting that localized protein translation is not required for the mRNA targeting to the MB.

Next, we set out to determine if ribosomal components that are needed for translation are also present at the MB. Initially, we analyzed three different proteomic datasets of purified MBsomes using The Database for Annotation, Visualization, and Integrated Discovery (DAVID)^8, 23, 24^. Consistent with the possibility of localized MB-associated translation, there are numerous ribosomal and translation initiation factor proteins present in the MB proteome (Figure 3C). Not only were there ribosomal and translation initiation factor proteins present in the proteomic datasets, but they were also highly enriched according to the DAVID analysis enrichment score (Figure 3C). Finally, upon staining HeLa cells with antibodies against Ribosomal Protein L3 (RPL3), we observed the presence of RPL3 in the MB and intracellular bridge (marked by anti-Acetylated-Tubulin antibodies) (Figure 3D-F). While this does not prove that local translation is occurring, it supports the idea that the ribosomal translation machinery is present at the MB and can potentially perform active translation of MB-associated mRNAs.

### Midbodies are sites for active translation

While the presence of ribosomal machinery at the MB supports the idea of local MB-associated protein translation, it is yet to be shown that active translation actually occurs at the MB and intracellular bridge during late telophase. To test this possibility, we used puromycin as a marker for active translation. Puromycin inhibits translation by causing the formation of a puromycylated nascent peptide chain and leading to dissociation of ribosomes and premature chain release. Importantly, these puromycylated peptides accumulate at sites of active translation and can be detected using anti-puromycin antibodies. To test whether translation occurs at the MB, we treated HeLa cells for 10 min with either DMSO or puromycin. We found that an anti-puromycin signal in MBs can be observed in puromycin-treated cells (Figure 4A-D), and, importantly, pre-treatment of cells with cycloheximide (Figure 4A) blocked the anti-puromycin signal (Figure 4A-D). Cycloheximide functions by interfering with the translocation step in protein synthesis, thus preventing ribosomes from incorporating puromycin in nascent peptides. Finally, anti-CHMP4B and anti-puromycin signals co-localized at the MB and intercellular bridge (Figure 4E-H). The presence of anti-puromycin signal in the MB and intracellular bridge is consistent with our hypothesis that MB-enriched mRNAs are locally translated during telophase. However, it is important to note that our experiments do not rule out the possibility that puromycylated nascent peptides could be generated in the cell body and then diffuse into the MB. Considering that MBs are diffusion limited structures (due to high cross-linking of MB microtubules) and short treatment time (only 10 min) we consider that unlikely. However, further experiments will be needed to fully demonstrate that CHMP4B is translated at the MB (also see Figure 8).

**Figure 4.**
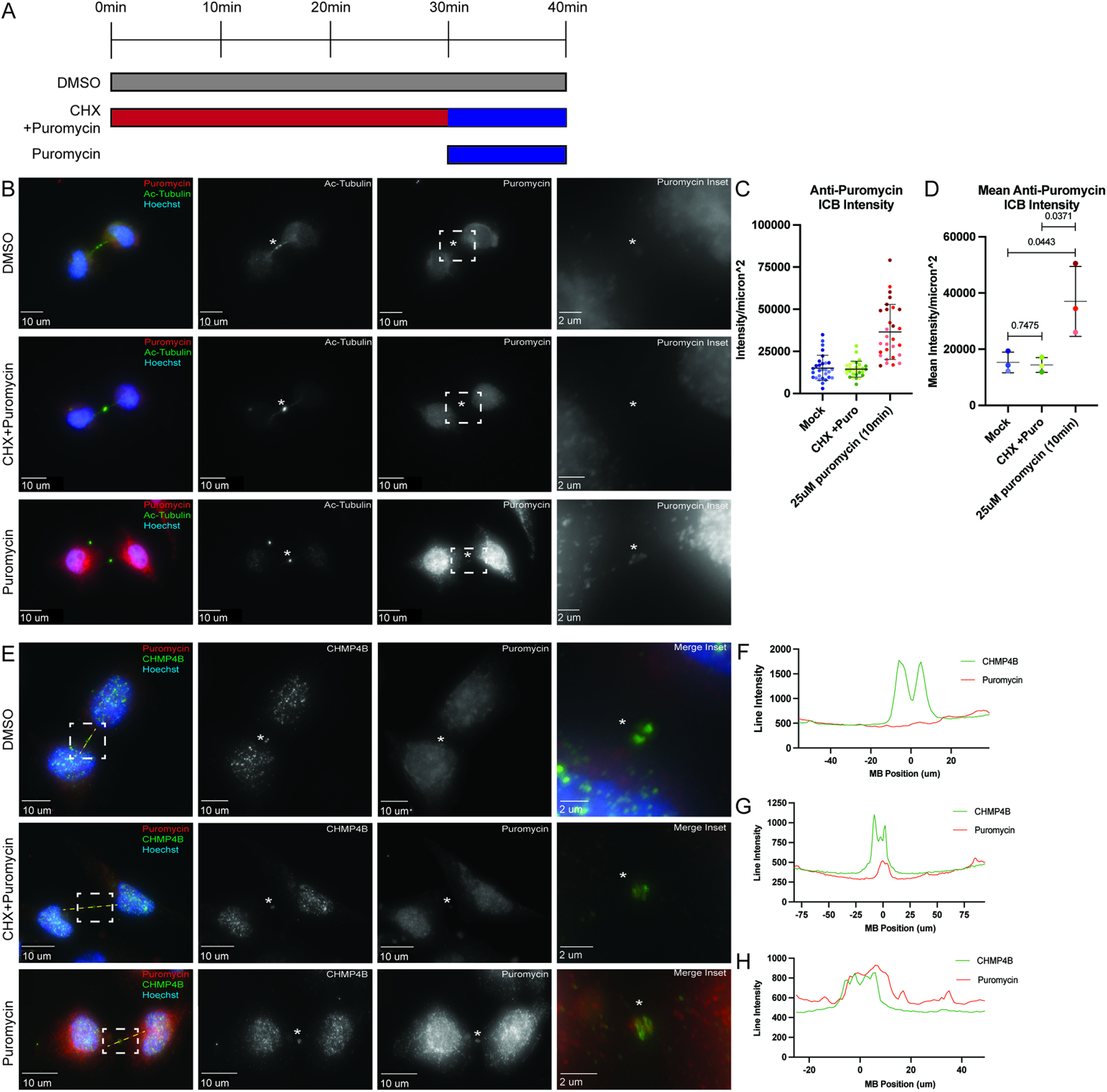
**Active protein translation occurs in the midbody.** (A) Schematic representing the 3 treatment conditions and times used for the experiments represented in panels (B-E). (B) HeLa cells were treated with either DMSO, CHX+Puromycin, or Puromycin and subjected to immunostaining with anti-acetylated-tubulin and anti-puromycin antibodies. The asterisks mark the MB, and the white dashed square represents the region of the image used for the inset. (C-D) 3i imaging software was used to measure the anti-puromycin intensity/micron^2^ in the intercellular bridge of 60 cells from 3 separate experiments (C; each dot represents one cell). Statistical analysis was done on means and standard deviations derived from the 3 individual experiments (D). Each experiment is represented by a different shade of color and is the mean derived from all cells analyzed in that particular experiment. Statistical analysis is represented with a p-value. (E) HeLa cells were treated with either DMSO, CHX+Puromycin, or Puromycin and subjected to immunostaining with anti-puromycin (to mark the puromycylated peptides) and anti-CHMP4B antibodies. The line used for the line intensity graphs is marked in yellow and the asterisks mark the MB. The white dashed square represents the region of the image used for the inset. (F-H) Line intensity graphs representing the intensity of Puromycin (red) and CHMP4B (green) signals after DMSO (F), CHX+Puromycin (G), or Puromycin (H) treatments.

### Active translation is required for the accumulation of CHMP4B at the MB during late telophase

Since our data suggest that local translation may occur in the MB, and that the anti-puromycin signal co-localizes with CHMP4B, we hypothesized that inhibiting protein translation at the MB should affect CHMP4B accumulation at the MB during late telophase. To test this hypothesis, we briefly (1 hour) treated HeLa cells with puromycin to block translation and analyzed endogenous CHMP4B levels at the MB by immunofluorescence microscopy. Consistent with our hypothesis, puromycin-treatment decreased CHMP4B intensity in the MB (Figure 5A-C). CHMP4B levels were also slightly reduced in the cell body, although to a much lesser (and statistically not significant) extent than seen in the MB (Figure 5A, D, and E).

**Figure 5.**
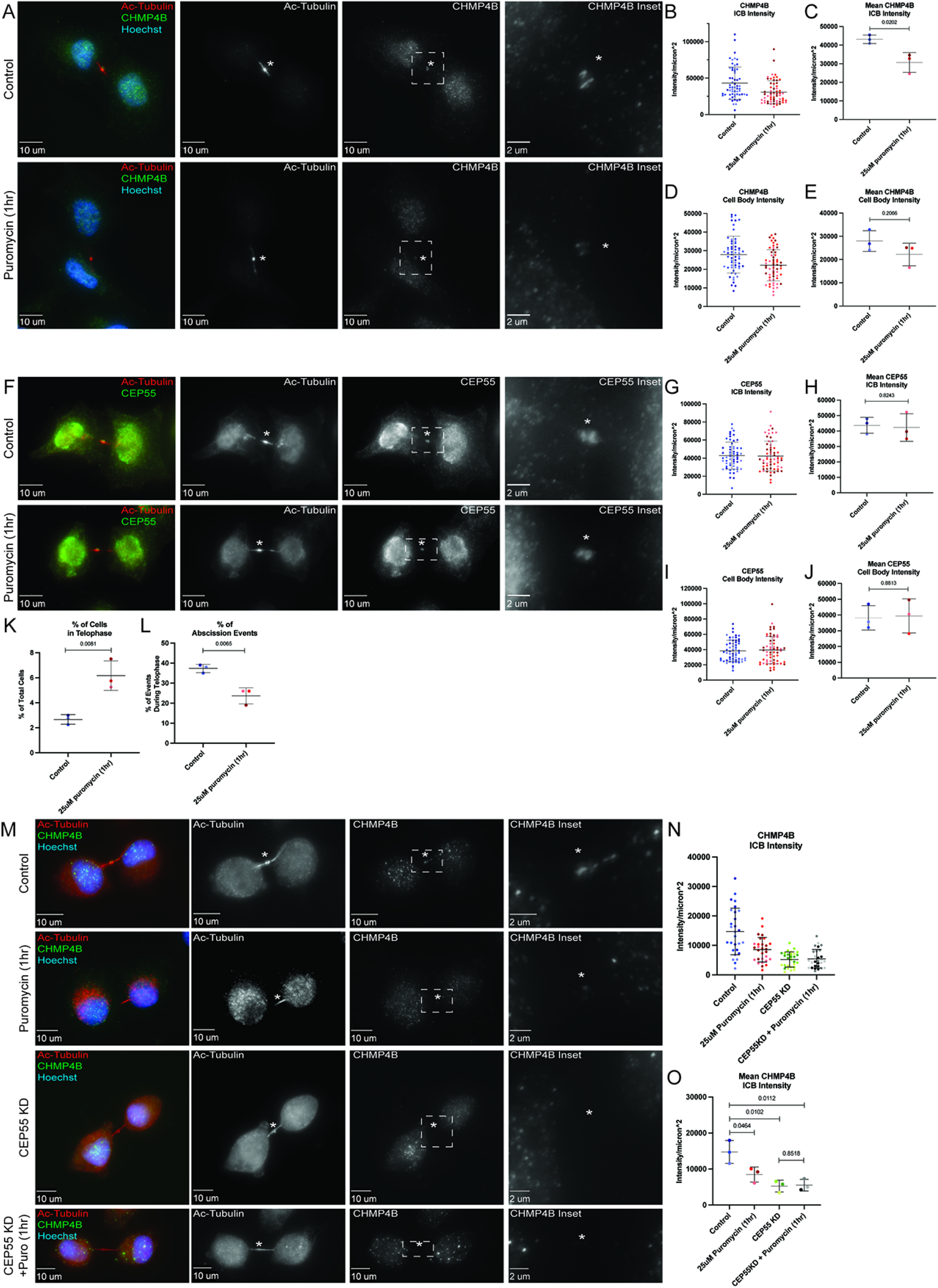
**Inhibition of translation inhibits CHMP4B accumulation at the midbody.** (A) HeLa cells were either untreated or treated with puromycin for 1 h and subjected to immunostaining with anti-acetylated-tubulin and anti-CHMP4B antibodies. The asterisks mark the MB, and the white dashed square represents the region of the image used for the inset. (B-C) Shown data represents CHMP4B intensity/micron^2^ in the intercellular bridge of 60 cells from 3 separate experiments. Panel B shows distributions derived from individual cells (each dot represents a single cell). Statistical analysis in panel C was done on the means and standard deviations derived from the 3 individual experiments where experimental means were calculated by averaging values from all the cells from each experiment. Each experiment is represented by a different shade of color. Statistical analysis is represented with a p-value. (D-E) Shown data represents CHMP4B intensity/micron^2^ in the cell body of 60 cells from 3 separate experiments. Panel D shows distributions derived from individual cells (each dot represents a single cell). Statistical analysis in panel E was done on the means, and standard deviations were derived from the 3 individual experiments where experimental means were calculated by averaging values from all the cells from each experiment. Each experiment is represented by a different shade of color. Statistical analysis is represented with a p-value. (F) HeLa cells were either untreated or treated with puromycin for 1 h and subjected to immunostaining with anti-acetylated-tubulin and anti-CEP55 antibodies. The asterisks mark the MB, and the white dashed square represents the region of the image used for the inset. (G-H) Shown data represents CEP55 intensity/micron^2^ in the intercellular bridge of 60 cells from 3 separate experiments. Panel G shows distributions derived from individual cells (each dot represents a single cell). Statistical analysis in panel H was done on the means, and standard deviations were derived from the 3 individual experiments where experimental means were calculated by averaging values from all the cells from each experiment. Each experiment is represented by a different shade of color. Statistical analysis is represented with a p-value. (I-J) Shown data represents CEP55 intensity/micron^2^ in the cell body of 60 cells from 3 separate experiments. Panel I show the distributions derived from individual cells (each dot represents a single cell). Statistical analysis in panel J was done on the means, and standard deviations were derived from the 3 individual experiments where experimental means were calculated by averaging values from all the cells from each experiment. Each experiment is represented by a different shade of color. Statistical analysis is represented with a p-value. (K) HeLa cells after puromycin treatment were fixed and stained with anti-acetylated tubulin antibodies. The percentage of cells in telophase was then counted. Shown data are the means and standard deviations derived from three independent experiments. Statistical analysis is represented with a p-value. (L) HeLa cells after puromycin treatment were fixed and stained with anti-acetylated tubulin antibodies. The percentage of cells that just completed abscission was then counted. Shown data are the means and standard deviations derived from three independent experiments. Statistical analysis is represented with a p-value. (M) HeLa cells were either untreated, treated with puromycin, CEP55 siRNA, or CEP55 siRNA plus puromycin and subjected to immunostaining anti-acetylated-tubulin and anti-CHMP4B antibodies. The asterisks mark the MB, and the white dashed square represents the region of the image used for the inset. (N-O) Shown data represents CHMP4B intensity/micron^2^ in intracellular of 60 cells from 3 separate experiments. Panel N shows distributions derived from individual cells (each dot represents a single cell). Statistical analysis in panel O was done on the means, and standard deviations were derived from the 3 individual experiments where experimental means were calculated by averaging values from all the cells from each experiment. Each experiment is represented by a different shade of color. Statistical analysis is represented with a p-value.

Puromycin blocks all protein translation. Consequently, even a short puromycin treatment will impact many proteins, thus, it is possible that inhibition of translation also affects other MB proteins. To further bolster this idea, we chose to look at CEP55, a protein that is recruited to the MB during its formation and is required for MB-accumulation of TSG101 and ESCRT-III complex (including CHMP4B) ^3, 4, 6, 9, 10, 16, 17^. Since CEP55 mRNA was not found to be enriched in our RNAseq analysis (Figure 2B), it presumably does not require MB-associated translation for its targeting. Consistent with this hypothesis, puromycin treatment did not have any effect on CEP55 intensity in the MB or cell body (Figure 5F-J).

The experiments described above used non-synchronized cells. Consequently, we cannot discount the possibility that 1 hour puromycin treatment affected CHMP4B mRNA translation during metaphase and anaphase before the formation of the MB. If that was the case, the differences we observed in CHMP4B accumulation at the MB would be the result of translation inhibition in the cell body (before telophase) rather than MB. To discount this possibility we next synchronized HeLa cells stably expressing GFP-MKLP1 using a thymidine/nocodazole block (Supplemental Figure 3A). Cells in metaphase were then washed to release from the block and incubated in serum-supplemented media for 90 minutes at 37°C followed by fixation and staining with (Supplemental Figure 3A). We have previously shown that within this 90-minute incubation cells progress from metaphase to telophase^45^, and, consistent with our previous work, we observed that over 90% of cells were in late telophase with clearly visible CHMP4B present at the MB (Supplemental Figure 3B). Importantly, the addition of puromycin for the last 30 minutes of the 90-minute incubation significantly decreased accumulation of endogenous CHMP4B at the MB (Supplemental Figure 3B-C).

If localized translation of CHMP4B, and presumably other members of the ESCRT-III complex, is required for their accumulation at the MB, we would expect that inhibition of translation during telophase would affect the cells’ ability to undergo abscission, by arresting them at late telophase. To test this hypothesis, we counted the number of cells in telophase and found that 1 hour of treatment with puromycin significantly increased the number of cells in telophase (Figure 5K), while decreasing the number of cells that have just undergone abscission (Figure 5L; Supplemental Figure 4A-D), which are all indications of defects in cell division.

To further test whether puromycin can delay cell abscission we next imaged telophase cells expressing GFP-MKLP1 for 90 minutes by time-lapse microscopy (Supplemental Figure 4E-F). In all cases we selected cells that were already in telophase, as indicated by the flattened cell, round nucleus, and presence of a GFP-MKLP1 labeled MB. Consistent with several previous studies, all cells (5 out of 5 imaged) in telophase successfully divided within 75 minutes (Supplemental Figure 4E; asterisk marks the MB; arrow points to abscission site). In contrast, none of the telophase cells (0 out of 5 imaged) completed abscission within 90 minutes if puromycin was added at the beginning of time-lapse analysis.

While the previous experiments support the idea that local translation of CHMP4B is at least partially responsible for the accumulation of CHMP4B during telophase, it does not exclude other mechanisms for CHMP4B accumulation at the MB. Since it has been proposed that CEP55 is the key initial recruiter of abscission-related proteins in mammalian cells, it is plausible that both pathways are required for efficient recruitment of ESCRT-III during abscission. To further test this idea, we used puromycin, CEP55 siRNA, or CEP55 siRNA/puromycin treatments and tested their effect on CHMP4B accumulation in the MB. As we already have shown, puromycin treatment alone caused a significant reduction in CHMP4B intensity at the MB (Figure 5M-O). Similarly, consistent with previous reports, CEP55 knock-down (Supplemental Figure 2B) also decreased CHMP4B intensity at the MB (Figure 5M-O). This decrease was not due to a decrease in *CHMP4B* mRNA accumulation at the MB (Supplemental Figure 2E) but likely due to lack of CEP55-dependent tethering of Alix and TSG101 at the MB (Figure 2D). Surprisingly, a combination of CEP55 KD and puromycin treatment did not further reduce CHMP4B intensity at the MB, suggesting that local MB-associated translation and CEP55 may work together to allow efficient accumulation (by localized translation) and retention (by tethering CHMP4B at the MB) of CHMP4B during late telophase.

### Short puromycin wash-out recovers CHMP4B levels at the MB but not cell body

While our previous data shows that MB-localized protein translation contributes to CHMP4B accumulation at the MB, the data are somewhat difficult to interpret because puromycin also resulted in a slight decrease of CHMP4B levels in the cell body. Thus, we could not fully differentiate between two possibilities: (1) CHMP4B targeting relies on MB-associated translation during telophase; (2) CHMP4B is translated at the cell body and then simply diffuses back to the MB. To help differentiate between those two possibilities we performed a puromycin wash-out experiment (Figure 6). Briefly, we treated cells with puromycin for 1 hour, followed by a quick puromycin wash-out and incubation of cells for 5 or 10 minutes, before analyzing the levels of CHMP4B in the MB and the cell body. As we already have shown, puromycin treatment resulted in decreased CHMP4B levels in both the MB and the cell body (Figure 6A-E). Importantly, after puromycin wash-out, CHMP4B levels at the MB started to increase within 5 min, with a complete recovery after 10min (Figure 6A, B-C). On the contrary, CHMP4B intensity in the cell body did not recover even after 10min (Figure 6A, D-E), suggesting that CHMP4B accumulation at the MB is primarily driven by localized translation rather than by diffusion of CHMP4B from the cell body.

**Figure 6.**
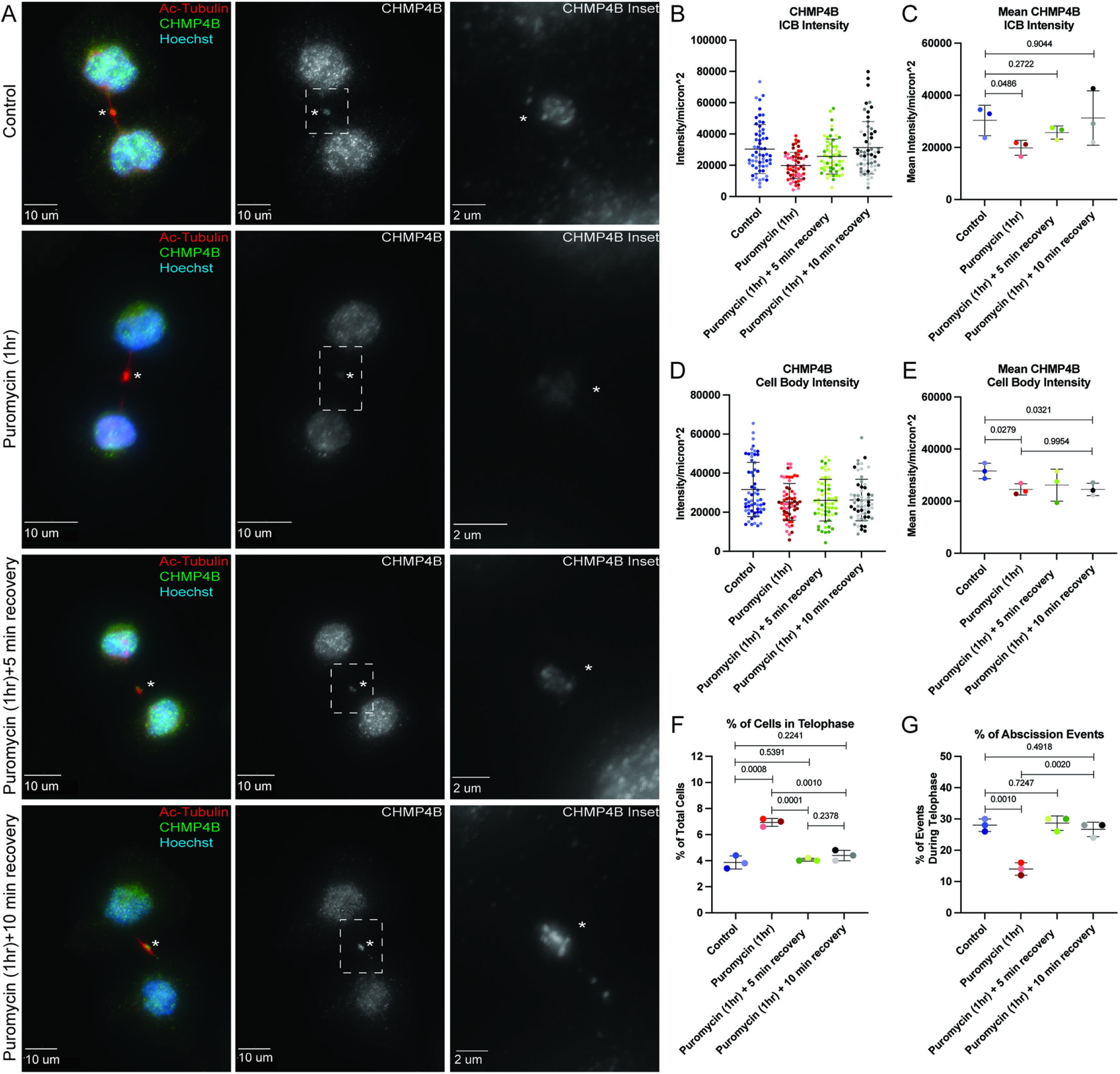
**Puromycin wash-out leads to fast recovery of CHMP4B levels in MB.** (A) HeLa cells were either untreated or treated for 1 h with puromycin. Cells were then washed and incubated with complete media for either 5 min or 10 min. Cells were then fixed and subjected to immunostaining with anti-acetylated-tubulin and anti-CHMP4B antibodies. The asterisks mark the MB, and the white dashed square represents the region of the image used for the inset. (B-C) Shown data represents CHMP4B intensity/micron^2^ in the intercellular bridge of 60 cells from 3 separate experiments. Panel B shows distributions derived from individual cells (each dot represents a single cell). Statistical analysis in panel C was done on the means, and standard deviations were derived from the 3 individual experiments where experimental means were calculated by averaging values from all the cells from each experiment. Each experiment is represented by a different shade of color. Statistical analysis is represented with a p-value. (D-E) Shown data represents CHMP4B intensity/micron^2^ in the cell body of 60 cells from 3 separate experiments. Panel D shows distributions derived from individual cells (each dot represents a single cell). Statistical analysis in panel E was done on the means, and standard deviations were derived from the 3 individual experiments where experimental means were calculated by averaging values from all the cells from each experiment. Each experiment is represented by a different shade of color. Statistical analysis is represented with a p-value. (F-G) HeLa cells that were untreated or treated for 1 h with puromycin. Cells were then washed and incubated with media for an additional 5 or 10 minutes. Cells were then fixed and stained with anti-acetylated-tubulin antibodies. The percentage of cells in either telophase (F) or just after abscission (G) was then counted. Shown data are means and standard deviations derived from three independent experiments. Statistical analysis is represented with a p-value.

Since CHMP4B levels rapidly recover at the MB after the puromycin wash-out, this should also lead to a rapid recovery of the cell’s ability to undergo abscission. Consistent with this idea, 5 minutes after the puromycin wash-out, cells fully recovered their ability to undergo abscission, as determined by a decrease in number of cells in telophase and increase in number of cells that have just undergone abscission (Figure 6F). All this data suggests that CHMP4B accumulation at the MB depends on local translation rather than diffusion from the cell body, and that MB-associated translation of CHMP4B is required for successful completion of the abscission.

### The 3′ UTR of CHMP4B mRNA enhances CHMP4B protein accumulation at the MB

Up to this point, we have been able to show that CHMP4B mRNA is present in the MB, local translation may occur at the MB, and that CHMP4B accumulation at the MB is at least partially regulated by local translation. The next step was to determine how the *CHMP4B* mRNA is targeted to the MB. RNA trafficking is often driven by cis-regulatory localization elements (LEs) that are commonly found in the 5’- or 3′ UTRs^28^. To that end, we generated clones containing the 5’- and/or 3′ UTR of *CHMP4B* fused to RFP-*CHMP4B* cDNA. One major issue with transient expression is the high variation in expression levels in different cells hindering our ability to tell whether differences in MB targeting of various *CHMP4B* constructs are due to UTRs or due to the differences in expression levels. To minimize this issue, we used a HeLa LoxP recombination system^29^. Briefly, this system uses *cre* recombination to allow for single copy, site-specific recombination of our constructs into the HeLa genome^29^. This allowed us to obtain virtually homogenous cell pools containing various doxycycline-inducible *CHMP4B* constructs. Importantly, each cell in these pools will have one copy of the construct that is inserted at the same genomic locus. Upon making cell lines containing various *CHMP4B* constructs, we first used immunofluorescence microscopy, western blotting, and qPCR to confirm that doxycycline does induce the expression of these constructs to the approximately the same level (Supplemental Figure 4A-G). Interestingly, we found that the presence of *CHMP4B* 3′ UTR without the 5′ UTR slightly deceased *CHMP4B* mRNA levels, raising the possibility that the 3′ UTR of CHMP4B may also regulate mRNA stability (Supplemental Figure 4B and E). Additionally, please note that the insertion of the 5′ UTR generated an alternative start codon that was in frame with the RFP, resulting in two CHMP4B bands on Western blots (Supplemental Figure 4F).

To determine the role of UTRs in CHMP4B targeting, we first tested whether having the 5’- and/or 3′ UTR of *CHMP4B* resulted in more efficient RFP-CHMP4B accumulation in the MB. To that end, we calculated the ratio of RFP-CHMP4B protein intensity at the MB and cell body. As shown in the figure 7A-C, the presence of the 3′ UTR significantly enhanced RFP-CHMP4B enrichment at the MB as compared to RFP-CHMP4B alone (Figure 7A-C). Interestingly, the 5′ UTR alone did not enhance RFP-CHMP4B targeting to the MB, nor did it enhance the effect of 3′ UTR (Figure 7A-C). All this data suggests that the 3′ UTR plays an important role in mediating CHMP4B protein accumulation in the MB during telophase, presumably due to targeting and localized translation of RFP-CHMP4B mRNA. Consistent with previous reports, RFP-CHMP4B without UTRs is still able to accumulate in the MB, although at much lower levels as compared to RFP-CHMP4B-3′UTR, potentially showing that there are multiple mechanisms, such as CEP55 binding, that drives ESCRT-III targeting during abscission.

**Figure 7.**
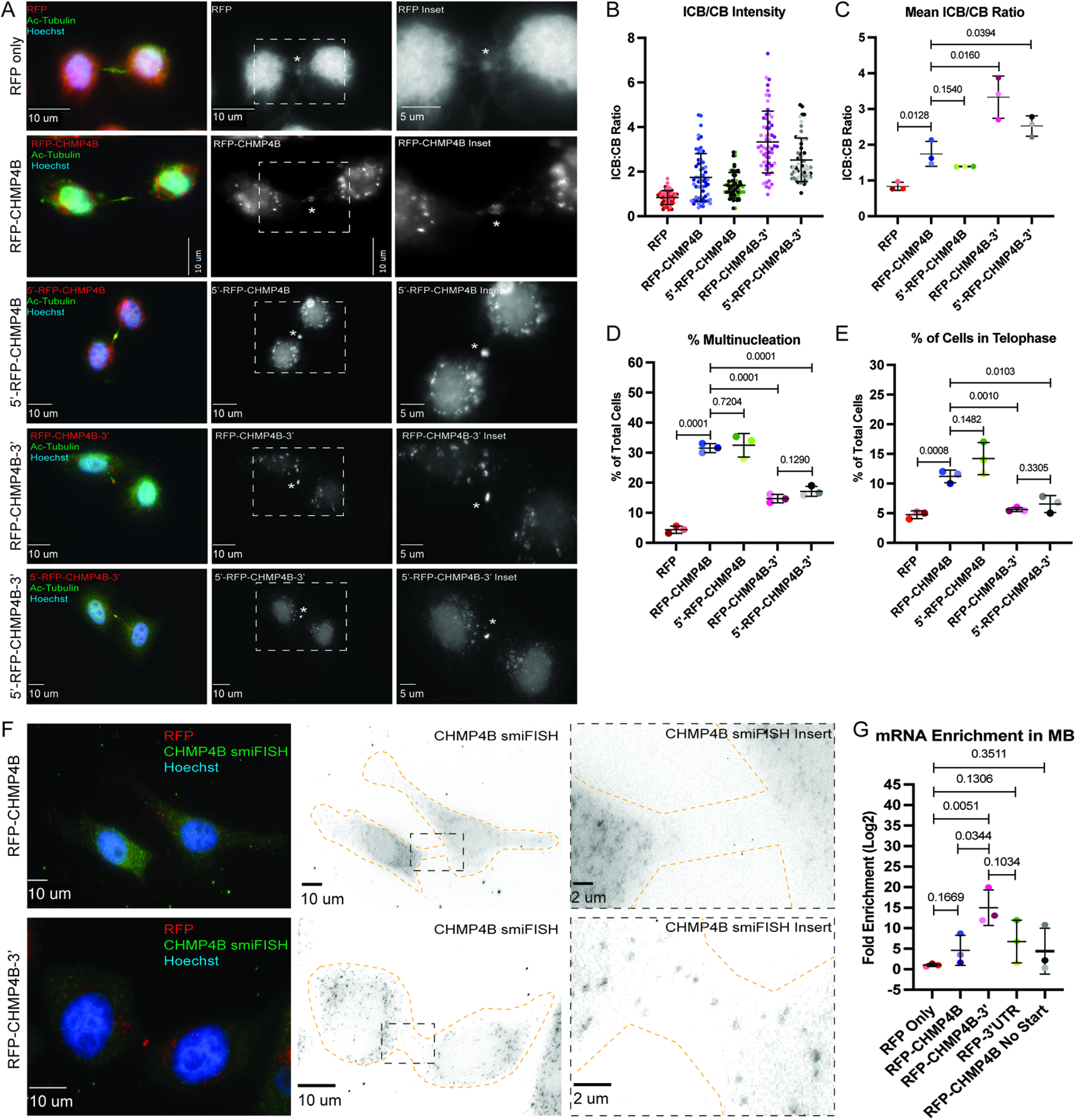
**The 3′ UTR of CHMP4B is important for protein accumulation and mRNA translocation.** (A) HeLa cells expressing various RFP-tagged dox-inducible constructs were incubated with 2μg/mL doxycycline for 48 h and then subjected to immunostaining with anti-acetylated-tubulin. The asterisks mark the MB, and the white dashed square represents the region of the image used for the inset. (B-C) Ratio between RFP fluorescence in the intracellular bridge and cell body was calculated from 60 randomly chosen cells in telophase. Panel B shows distributions derived from individual cells (each dot represents a single cell). Statistical analysis in panel C was done on the means, and standard deviations were derived from the 3 individual experiments where experimental means were calculated by averaging values from all the cells from each experiment. Each experiment is represented by a different shade of color. Statistical analysis is represented with a p-value. (D) HeLa cells expressing various RFP-tagged dox-inducible constructs were incubated with 2μg/mL doxycycline for 48 h and then subjected to immunostaining with anti-acetylated-tubulin. The percentage of cells with multinucleation was then counted. Shown data are the means and standard deviations derived from three independent experiments. Statistical analysis is represented with a p-value. (E) HeLa cells expressing various RFP-tagged dox-inducible constructs were incubated with 2μg/mL doxycycline for 48 h and then subjected to immunostaining with anti-acetylated-tubulin. The percentage of cells in telophase was then counted. Shown data are the means and standard deviations derived from three independent experiments. Statistical analysis is represented with a p-value. (F) HeLa cells expressing dox-inducible RFP-CHMP4B or RFP-CHMP4B-3’UTR constructs were incubated with 2μg/mL doxycycline for 48 h and subjected to staining with smFISH probes against CHMP4B. The cell outline is marked in yellow with the asterisks marking the MB. The black dashed square represents the region of the image used for the inset. (G) Whole cell or MB-associated mRNA was isolated from these cell lines expressing various RFP-tagged constructs and subject to RT-qPCR analysis using RFP or GAPDH (control) specific primers. RFP mRNA levels were then normalized against GAPDH mRNA and expressed as a ratio between MB and whole cell RFP mRNA levels. Statistical analysis is represented with a p-value and was done on the means and standard deviations derived from three independent experiments.

It has been previously reported that over-expression of exogenous tagged ESCRT-III members inhibits abscission^13^. Consistent with those studies, we also see an increase in multinucleation and the number of telophase-arrested cells in the cells overexpressing RFP-CHMP4B (Figure 7D-E). While the cause of this inhibition remains to be fully understood, it has been suggested that CHMP4B overexpression may lead to accumulation of cytosolic ESCRT-III aggregates, thus, sequestering the rest of the ESCRT-III subunits away from the MB. We wondered whether the inhibitory effect of exogenous CHMP4B may be due to the fact that it’s mRNA lacks localization elements mediating targeting of CHMP4B to the MB. Consistent with this hypothesis, the effect of over-expression on multinucleation or telophase arrest was reversed in cell lines expressing RFP-CHMP4B constructs containing 3′ UTR (Figure 7D-E).

### The 3′ UTR of CHMP4B mRNA is required for its targeting to the MB

Since the 3′ UTR of *CHMP4B* RNA is important for CHMP4B protein accumulation at the MB during abscission, it might be expected that the 3′ UTR is also important for *CHMP4B* mRNA targeting to the MB. To determine that, we first analyzed *RFP-CHMP4B* mRNA localization using smiFISH probes against *CHMP4B*. Consistent with the requirement of the 3′ UTR for RFP-CHMP4B accumulation, we observed smiFISH signal in the MB of cells only expressing the RFP-CHMP4B-3′UTR construct (Figure 7F). One drawback of using smiFISH in the MB is that it is only semi-quantifiable due to not knowing how efficiently the probes are able to penetrate the protein-rich region. To better quantify the difference between *RFP-CHMP4B* mRNA targeting to the MB, we purified MBsomes from cell lines expressing various constructs and analyzed mRNA levels by performing RT-qPCR on purified MBsomes and then compared them to whole cell mRNA levels. To measure only exogenous *CHMP4B* RNA, we used primers designed to target the *RFP* sequence. As shown in Figure 7G, the presence of the 3′ UTR was required for efficient *RFP-CHMP4B* mRNA targeting to the MB, as compared to the amount of *RFP* mRNA alone or *RFP-CHMP4B* mRNA without 3′ UTRs (Figure 7G). This data suggests that the 3′ UTR is required for efficient *RFP-CHMP4B* mRNA targeting to the MB. Interestingly, *RFP-CHMP4B* RNA without UTRs was also slightly enriched at the MB as compared to *RFP* RNA alone. This slight mRNA enrichment could be due to co-translational *RFP-CHMP4B* mRNA accumulation at the MB. To test this possibility, we removed the start codon and generated a HeLa cell line expressing doxycycline-inducible RFP-CHMP4B-NoStart construct (Supplemental Figure 4C and E) and tested whether this mRNA also accumulates at the MBs. As shown in Figure 7E, removal of the start codon did not affect the ability of *RFP-CHMP4B* mRNA to accumulate at the MB, although it was clearly still much lower than *RFP-CHMP4B-3′UTR*. All this data suggests that the 3′ UTR is required for efficient CHMP4B mRNA targeting, although the coding sequence also appears to contribute to mRNA accumulation at the MB.

To test whether the 3′ UTR of CHMP4B is sufficient for the targeting of any mRNA to the MB, we generated a doxycycline inducible RFP-only construct that contained the 3′ UTR from *CHMP4B* (RFP-3’UTR; Supplemental Figure 4D and E) and tested the mRNA levels at the MB. As shown in the figure 7E, 3′ UTR of *CHMP4B* did slightly enhance the levels of *RFP* mRNA at the MB, although it did not reach the levels of *RFP-CHMP4B-3′UTR* mRNA. This suggests that while *CHMP4B* 3′ UTR is essential for mRNA targeting, it may require other targeting elements that are possibly present in the *CHMP4B* coding region or secondary structure of the coding region to get the full targeting of mRNA to the MB.

### The 3′ UTR of CHMP4B mRNA is required for its localized translation at the MB

Since the 3′ UTR is required for *CHMP4B* mRNA targeting to the MB, we wondered whether it may also be required for localized CHMP4B translation at the MB. If CHMP4B is locally translated at the MB, it would be expected to start accumulating at the MB during late telophase after the formation of the MB. Indeed, it has been previously shown that endogenous CHMP4B accumulates at the MB just before the abscission, while CEP55, TSG101, and Alix are known to be present at the MB from the very beginning of MB formation. To understand the timing and dynamics of RFP-CHMP4B accumulation at the MB we next performed a time-lapse analysis of cells stably expressing *RFP-CHMP4B-3′UTR* mRNA (Figure 8A). To minimize the possibility that RFP-CHMP4B diffuses from the cell body we chose cells in late telophase (as determined by round nuclei and flattened body) that were still connected with a long intracellular bridge (Figure 8A, MB marked by asterisk). As shown in the figure 8A, RFP-CHMP4B was initially absent in the MB (see time point 0 min), but then started accumulating with the MB over 10-30 minutes, the dynamics consistent with translation of CHMP4B at the MB during late telophase.

**Figure 8.**
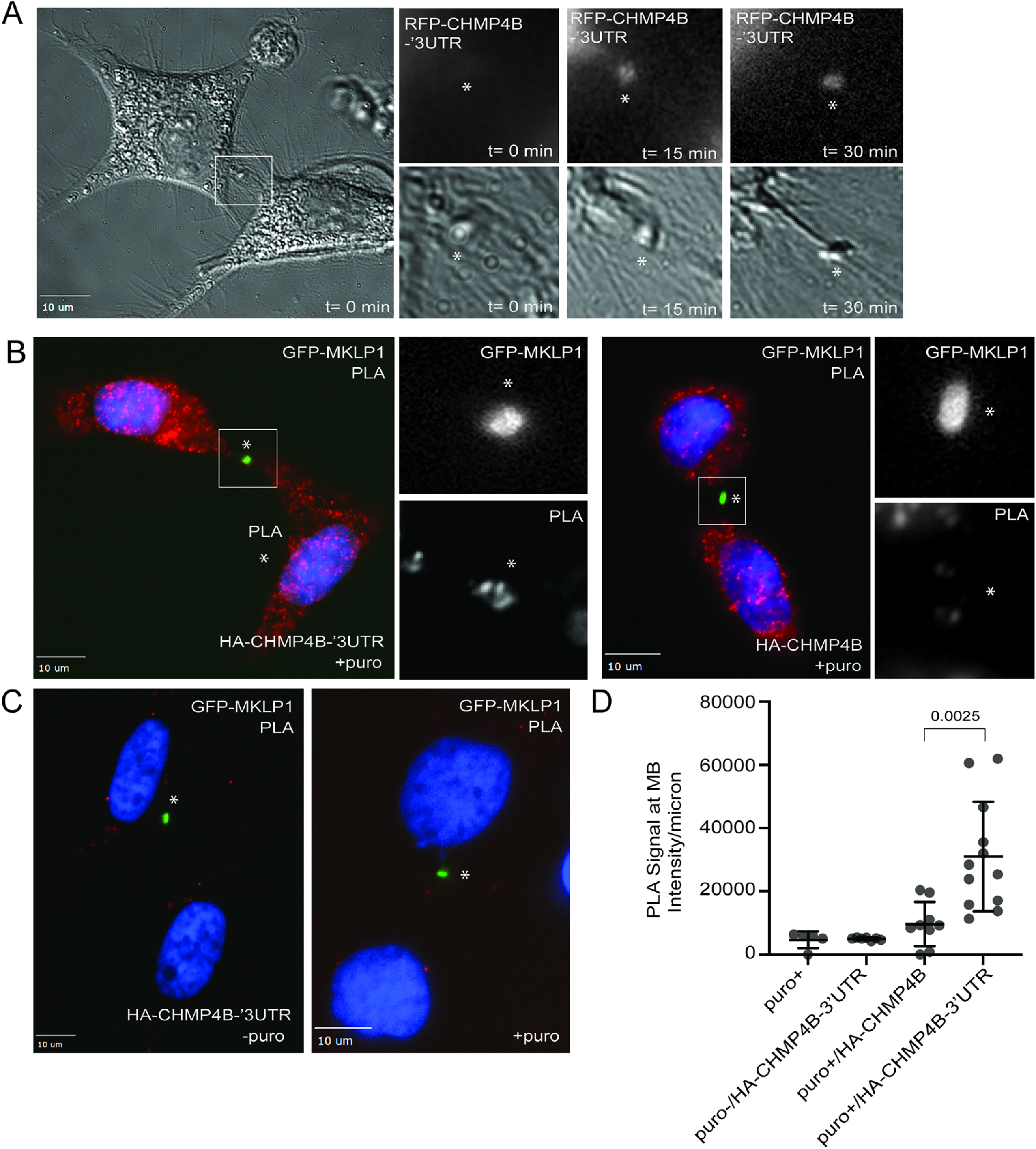
**The 3′ UTR is required for CHMP4B translation at the MB during telophase.** (A) HeLa cells were transfected with RFP-CHMP4B-3′UTR and plated on glass-bottom dishes. The accumulation of RFP-CHMP4B at the MB during telophase was then imaged using time-lapse microscopy for 120 minutes with 15 minute time-lapse. Asterisk marks the midbody (B) HeLa cells stably expressing GFP-MKLP1 were transfected with either HA-CHMP4B-3′UTR (images on the left) or HA-CHMP4B (images on the right). Cells were then fixed and incubated with anti-HA and anti-puromycin antibodies followed with proximity ligation assay (PLA). Asterisk marks the midbody. Boxes mark area shown in higher magnification insets. (C) Negative controls for PLA experiment shown in panel B. Cells in image on the left were not treated with puromycin but transfected with HA-CHMP4B-3′UTR. Cells in image on the right were treated with puromycin but not transfected with HA-CHMP4B-3′UTR. Asterisk marks the midbody. (D) Quantification of the levels of PLA signal in the midbody area. Data shown are the means and standard errors of mean. Each circle indicates the data derived from one dividing cell.

While the time-lapse analysis of cells expressing *RFP-CHMP4B-3′UTR* mRNA is consistent with the possibility that RFP-CHMP4B is locally translated at the MB, it does not directly demonstrate that CHMP4B is locally translated and that the 3′ UTR is required for this process. To more directly investigate that, we overexpressed an N-terminally HA-tagged CHMP4B construct with or without *CHMP4B*’s 3′ UTR. Cells were then treated for 10 minutes with puromycin, followed by fixation and incubation with anti-HA and anti-puromycin antibodies. We next analyzed fixed cells with proximity ligation assay (PLA). The idea of this experiment is that the accumulation of nascent peptides containing HA (at the N-terminus) and puromycin (more towards the C-terminus) will be an indication of local translation and can be detected by HA/puro PLA. Importantly, since the 3′ UTR of *CHMP4B* is required for *HA-CHMP4B* mRNA targeting to the MB, we should not detect PLA signal in cells overexpressing HA-CHMP4B lacking the 3′ UTR. This experiment also allowed us to test whether diffusion from the cell body of puromycylated nascent peptides is sufficient to generate PLA signal in the MB (also see experiments shown in Figure 4). As shown in the figure 8B-D, we could indeed detect HA/puro PLA signal in the cells expressing *HA-CHMP4B-3′UTR*. This signal was significantly diminished in cells expressing HA-CHMP4B without 3’UTR. Importantly, the PLA signal was essentially absent in cells not treated with puromycin or cells not expressing the HA-tagged CHMP4B construct (Figure 8C-D). Taken together, all these data demonstrate that 3′ UTR is required for *CHMP4B* mRNA targeting to the MB and is also needed for localized CHMP4B translation at the MB during late telophase.

### The MBsome transcriptome is similar to other localized transcriptomes associated with the plus-ends of microtubules

We next sought to identify specific *cis*-elements (LEs) within MBsome-enriched RNAs that regulated their trafficking to the MB. We recently identified RNA elements that were necessary and sufficient for transport to both the neurites of neuronal cells and the basal pole of epithelial cells^40^. Both of these subcellular locations are associated with the plus ends of microtubules, and we showed that these RNA elements directed RNAs to their destinations by targeting them to microtubule plus ends. Given that MBs are associated with the plus-ends of microtubules emanating from each daughter cell, we reasoned that perhaps the same regulatory elements and mechanisms that drive RNAs to neurites and the basal pole of epithelial cells would also drive mRNAs to MBs.

If this were true, we would expect the same sets of mRNAs to be enriched at all three subcellular locations. To test this, we asked whether our MBsome-enriched RNAs were more neurite-enriched than other RNAs. For this comparison, we used a previously published dataset in which mouse neuronal cells were mechanically fractionated into soma and neurite fractions followed by analysis of their RNA contents using RNAseq^46^. We compared the neurite-enrichments of the mouse orthologs of our MBsome-enriched RNAs to the neurite-enrichments of all other mouse RNAs. We found that the mouse orthologs of the MBsome-enriched RNAs were significantly more neurite-enriched than all other RNAs, suggesting that neurites and MBsomes contain similar mRNA contents (Figure 9A).

**Figure 9.**
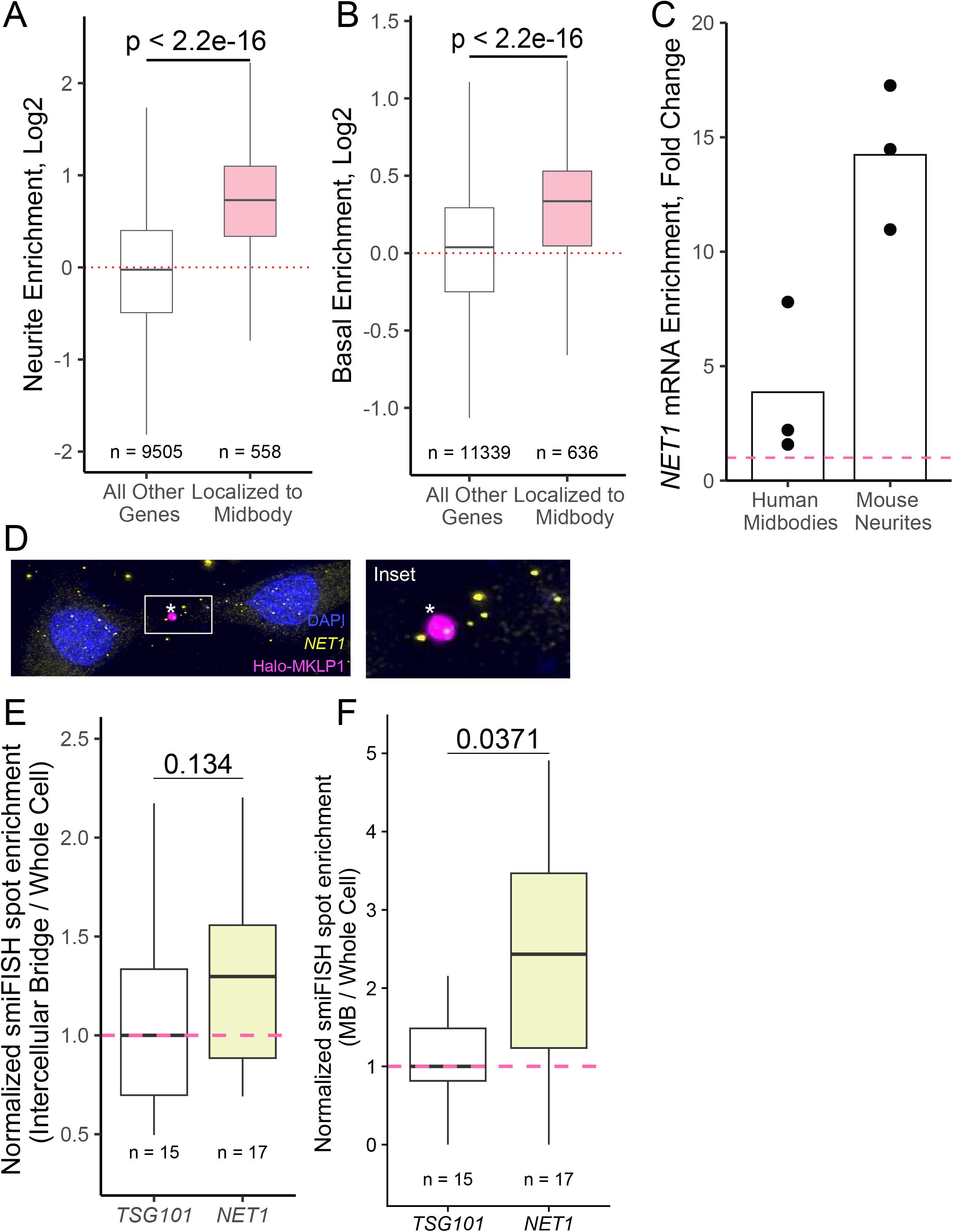
**Transcripts associated with the plus-ends of microtubules are enriched in the MB transcriptome.** (A) Comparison of MB-associated mRNA with RNAs enriched in neuronal projections of mouse neuronal cells. Genes were binned into MB-localized RNAs and non-MB-localized RNAs. Neurite RNA enrichment values (neurite RNA / soma RNA, log2) from previous published data were then calculated for the mouse orthologs of all genes, and the distributions of enrichment values were compared between bins. P values were calculated using a Wilcoxon rank-sum test. (B) As in A, genes were binned according to their MB RNA enrichment. Using a previous published comparison of the apical and basal compartments of epithelial cells, basal enrichment values (basal RNA / apical RNA, log2) were calculated for all genes, and the distributions of enrichment values were compared between bins. P values were calculated using a Wilcoxon rank-sum test. (C) *NET1* RNA enrichment in human midbodies compared to whole cells (left) and mouse neurites compared to cell bodies (right). (D) smiFISH for endogenous *NET1* mRNA localization (yellow) in cells stably expressing Halo-MKLP1 (pink). Midbody is marked with an asterisk. The box indicates the area shown in a higher magnification inset. (E) Quantification of *NET1* smiFISH. Transcript counts were quantified in the intercellular bridge and whole cell areas. For each cell, the number of counts in the intercellular bridge was normalized by the number of counts in the whole cell. The median ratio for this value across all cells for the control *TSG101* transcript was set to one. P values were calculated using a t-test. (F) As in E, except that transcript counts in the midbody and whole cell regions were compared. P values were calculated using a t-test.

Using data from human intestinal epithelial cells^40^, we then compared enrichments across the apicobasal axis for MBsome RNAs and all other detected RNAs. Again, we found that MBsome-enriched RNAs were significantly enriched at the basal pole of these cells compared to non-MBsome-enriched RNAs (Figure 9B). Taken together with the neurite RNA enrichments observed above, this indicates that neurites, the basal pole of epithelial cells, and MBsomes are enriched for similar sets of mRNAs, implying that similar mechanisms (e.g. plus-end directed transport along microtubules) may be populating their transcriptomes.

### Identification of 3′ UTR localization elements that are necessary and sufficient to target mRNA to the MB

Our previous studies that identified mechanisms behind RNA transport along microtubules focused on the *Net1* mRNA^40, 47^. These studies identified a GA-rich 260 nt region within the 3′ UTR of *NET1* that was both necessary and sufficient to direct RNA to the plus ends of microtubules in both neuronal and epithelial cells. We therefore wondered if the same element had similar abilities to direct RNA transport to the MB and MBsomes.

We started by asking if endogenous *NET1* mRNA was enriched in our MBsome RNA samples (Figure 1D) since our previous studies had found *NET1* mRNA enriched in the neurites^47^, and we observed *NET1* mRNA enriched at the basal pole of intestinal epithelial cells (Supplemental Figure 6A, B). We found that *NET1* mRNA was approximately 4-fold enriched in MBsomes relative to whole cells (Figure 9C). This observation was confirmed with smiFISH, showing *NET1* mRNA puncta in close proximity to MBs in dividing cells (Figure 9D). After quantifying the smiFISH results across many cells, we found that *NET1* mRNA was more likely to be enriched at both the intracellular bridge (Figure 9E) and the midbody (Figure 9F) compared to the control transcript *TSG101* (that is not enriched in MBs as measured by RNAseq and qPCR; see Figure 2).

We next asked if the same RNA *cis*-regulatory element within mouse *Net1* mRNA previously shown to promote RNA trafficking to neurites and the basal pole of epithelial cells also controlled RNA localization to MBs and MBsomes. To test this, we devised an RT-qPCR-based reporter assay that quantified the enrichment of designed reporter transcripts in MBsomes (Figure 10A). In this assay, Firefly and Renilla luciferases are co-expressed from a bidirectional promoter with the Renilla luciferase RNA serving as an unchanging internal control. RNA from MBsomes and whole cells was collected as described before, and the relative amounts of Firefly and Renilla luciferase transcripts in each fraction was determined using Taqman RT-qPCR. By comparing how the ratios of these transcripts vary between cell fractions depending on the 3′ UTR appended to the Firefly luciferase RNA, we are able to quantify the ability of the 3′ UTR to drive the Firefly luciferase transcript to MBsomes.

**Figure 10.**
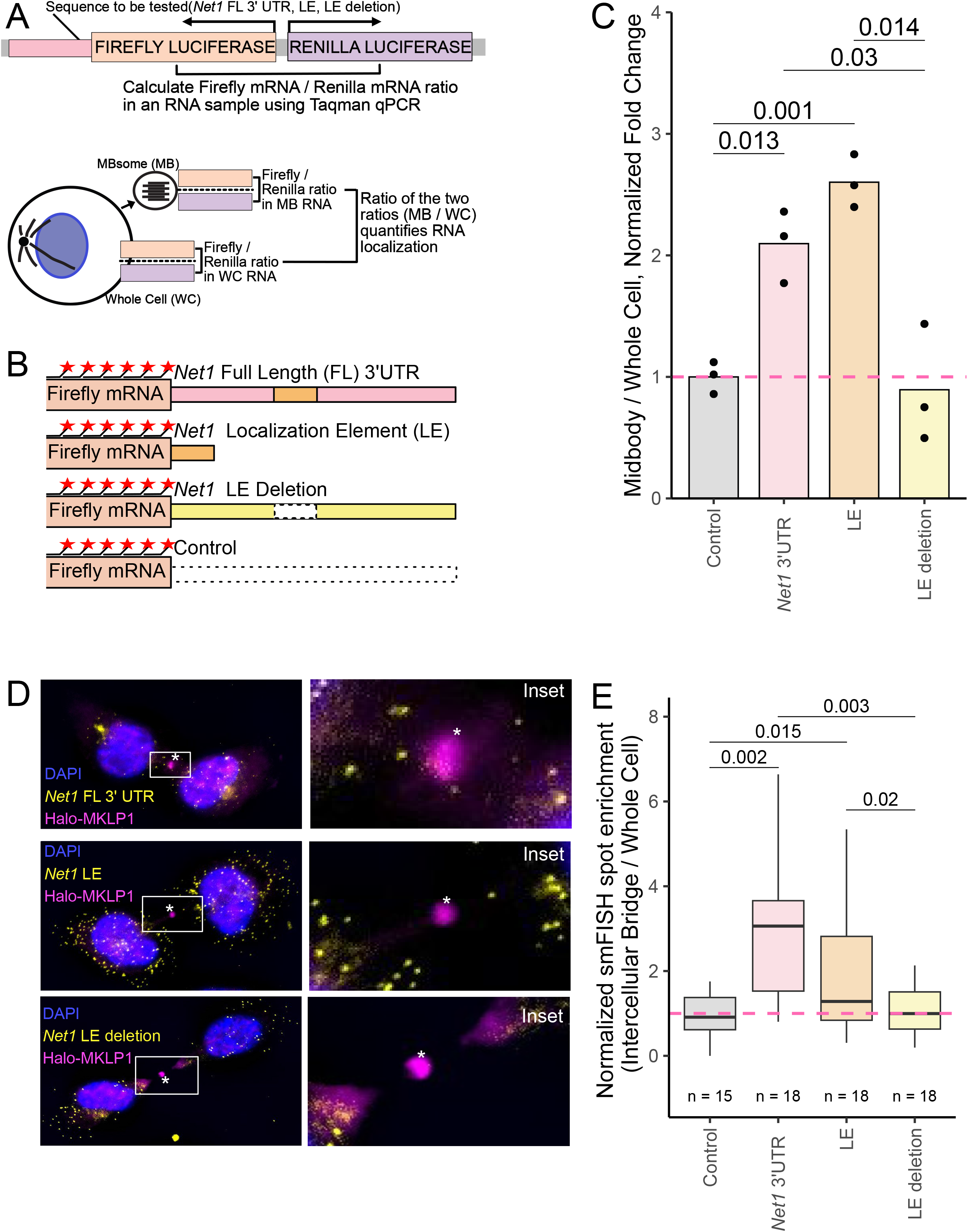
*NET1* 3′ UTR localization elements are necessary and sufficient to target mRNA to the MB (A) Schematic RT-qPCR-based reporter RNA approach for assaying MB RNA localization. Plasmids expressing Firefly and Renilla luciferases from a bidirectional reporter are integrated into the genome. Sequences to be tested for MB localization activity (*NET1* full length 3′ UTR, LE and LE deletion) are fused to the 3′ UTR of Firefly luciferase. Using Taqman qPCR, the ratio of Firefly to Renilla luciferase mRNA is measured in MB and whole cell samples. The ratio of these ratios (MB / WC) quantifies the MB-enrichment of the Firefly luciferase transcript. By asking how this value changes upon appending additional sequences to the Firefly luciferase RNA, the effects of these additional sequences on MB RNA enrichment can be tested. (B) Diagram of reporter constructs used to interrogate *NET1* localization activity in MBs. smFISH probes against the Firefly luciferase coding sequence are shown as red stars. (C) Localization of the reporter RNAs described in B to the MB as quantified using the strategy outlined in A. The value for the control reporter lacking additional sequence elements was set to one. P values were calculated using a t-test. (D) smFISH visualizing the reporter RNAs described in B (yellow) in cells stably expressing Halo-MKLP1 (pink). Midbody is marked with an asterisk. The box indicates the area shown in a higher magnification inset. (E) Quantification of MB-localized *Net1* UTR-containing reporter transcripts visualized in D. In all samples, reporter transcript counts in intercellular bridges and whole cells were quantified and the ratio of counts between the two locations is reported. These ratios were normalized by setting the value for the control reporter transcript lacking 3′ UTR additions to one. P values were calculated using a t-test.

We created four Firefly luciferase reporter RNAs (Figure 10B). These reporters contained 3′ UTRs consisting of the entire *Net1* 3′ UTR, the previously identified 260 nt element within the *Net1* UTR that was sufficient to drive RNA transport to neurites and the basal pole of epithelial cells, the entire *Net1* UTR lacking this element, and as a control, an unrelated plasmid-encoded UTR. These constructs were site-specifically incorporated into HeLa cells using *cre*/loxP recombination, and their expression was controlled using a doxycycline-on system.

We then measured the relative MBsome-enrichment of each of these constructs using the RT-qPCR strategy outlined in Figure 9A, setting the enrichment of the reporter with the control 3′ UTR to one. We found that the addition of the entire *Net1* UTR to the reporter construct increased its MBsome-enrichment by approximately 2-fold (Figure 10C). The reporter containing the previously identified 260 nt subset of the *Net1* UTR was similarly enriched in MBsomes, indicating that this *cis*-element is sufficient to drive RNAs to MBsomes. Deletion of this element completely abrogated the ability of the *Net1* 3′ UTR to regulate MBsome RNA localization, demonstrating that the element is also necessary for RNA transport to MBsomes.

Next, we confirmed these results using an orthogonal smFISH assay that visualized Firefly luciferase reporter RNAs *in situ*. The fluorescent probes used in this assay targeted the Firefly luciferase coding sequence that was shared across all reporter RNAs. As with the RT-qPCR approach, we found that the 260 nt *cis*-element within the *Net1* UTR was both necessary and sufficient for reporter RNA targeting to both MBs and the intercellular bridge of dividing cells (Figure 10D, E, Supplementary Figure 6C).

These results demonstrate that discrete RNA *cis-*regulatory elements control the RNA content of MBs and MBsomes and suggest that previously defined mechanisms that regulate microtubule-based RNA transport in other systems likely also act to populate the MBsome transcriptome^40^.

## Discussion

The midbody (MB) is a microtubule-rich structure that forms at the end of cytokinesis as the result of compacting central spindle microtubules during cleavage furrow ingression. Even though the MB was first identified over a hundred years ago, only in the last decade has it become apparent that the MB is not simply a byproduct of cell division but plays many important roles. It is now well-established that the MB functions as a scaffolding platform that recruits and activates many abscission regulating proteins, such as Aurora B kinase, MKLP1, and the ESCRT-III complex^16, 30–33^. Furthermore, recent studies have also shown that during cell division MBs can be released in the extracellular milieu where they are called MBsomes or Flemmingsomes^8, 23^. Uptake of these MBsomes by other cells can lead to changes in cell fate, differentiation, and proliferation capacity, presumably by mediating lateral transfer of specific signaling molecules^23, 24, 34^. However, what remains unclear is what these signaling molecules are and how they may affect cells after MBsome internalization. In this study we demonstrate that MBsomes contain a specific subset of mRNAs, many of them encoding proteins regulating cell division and differentiation. That raises a very intriguing possibility that delivery of specific mRNAs to the acceptor cells is what mediates post-mitotic MBsome functions, although further research will be needed to fully define the role of MBsome-dependent mRNA delivery.

Our study shows that MBs contain mRNAs encoding for several ESCRT-III components, namely CHMP2A, CHMP3, CHMP4B, and CHMP6. What makes this discovery especially exciting is that all these ESCRT-III complex proteins are known to co-oligomerize and drive the abscission step of cytokinesis. While the role of the ESCRT-III proteins in mediating abscission was first described over a decade ago^6, 9, 15^, how ESCRT-III subunits are delivered and targeted to the MB remains somewhat controversial. Until recently, it has been thought that TSG101 and the ESCRT-III complex accumulate at the MB by directly or indirectly binding to MB resident protein, CEP55^3, 4, 6, 9, 10, 16, 17^. However, recent studies have shown that depletion of CEP55 does not fully block ESCRT-III targeting to the MB, and CEP55 knock-out mice are able to grow and develop with limited defects^18^. Additionally, CEP55 is only present in vertebrates, raising a question of how the ESCRT-III may be targeted in other organisms, although CEP55 clearly contributes to ESCRT-III targeting in vertebrate cells. Finally, the MB and intercellular bridge are packed with dense and heavily crosslinked microtubule bundles^35, 36^, thus, creating a diffusion-limited environment. This raises the question about how large protein complexes, such as the ESCRT-III complexes, can rapidly diffuse via the intercellular bridge into the MB from the cytosol in a timely, efficient, and highly regulated manner. The discovery that ESCRT-III mRNAs are enriched at the MB suggests an intriguing possibility that regulated localized translation of ESCRT-III components contribute to accumulation of ESCRT-III at the MB during abscission. This hypothesis is supported by results in other systems in which precise RNA localization is critical for the successful association of the resulting translated protein with its proper interaction partners^48^.

Consistent with the hypothesis that localized protein translation contributes to ESCRT-III targeting to the MB, we show that the MB has all the components needed for translation. We also use anti-puromycin antibodies to demonstrate that active translation is occurring in the MB. Finally, we show that inhibition of protein translation during late telophase blocks completion of cytokinesis. Based on all these findings, we chose to further investigate one of the most important regulators of late-stage abscission, CHMP4B. There are several pieces of evidence suggesting that CHMP4B is being actively translated in the MB during telophase. First, puromycylated CHMP4B nascent peptides are present at the MB. Second, inhibition of protein translation during telophase leads to decrease in CHMP4B accumulation at the MB. Third, wash-out of the translation inhibitor puromycin results in a very fast recovery of CHMP4B levels at the MB, which could not be explained by simple diffusion of CHMP4B protein that was translated in the cell body. Importantly, the ability of the cell to complete abscission also recovers within a similar timescale. All of these data suggests that CHMP4B can be locally translated and that it is at least partially responsible for the accumulation and function of CHMP4B at the MB. Importantly, the mechanisms using of CEP55 and local translation to target ESCRT-III to the MB are not necessarily mutually exclusive. Indeed, both of these mechanisms appear to contribute to ESCRT-III localization in vertebrate cells suggesting that while localized translation may be required to accumulate ESCRT-III components within the MB, CEP55 is likely needed to tether and maintain their MB-enrichment during telophase.

Local translation of CHMP4B at the MB depends on the targeting of *CHMP4B* mRNA to the MB during telophase. What remains unclear is the molecular machinery governing mRNA delivery and accumulation at the MB. It has been previously shown that the subcellular targeting of mRNAs is usually driven by *cis*-regulatory elements that are commonly found in the 5’ or 3′ UTRs^28^. Consequently, we tested whether the 5’ and 3′ UTRs are driving the translocation of *CHMP4B* mRNA to the MB. Our data demonstrates that the 3′ UTR is required for translocation and accumulation of CHMP4B at the MB. To test whether the 3′ UTR is sufficient for mRNA accumulation at the MB, we added CHMP4B 3′ UTR to RFP and analyzed its ability to drive RFP mRNA accumulation at the MB. Interestingly, while we did see an increase in the levels of MB-associated *RFP* mRNA, the amount of *RFP-3′UTR* mRNA did not fully recapitulate the levels of *RFP-CHMP4B-3′UTR* mRNA, suggesting that the *CHMP4B* coding region may also play a role in *CHMP4B* mRNA targeting to the MB.

It has been previously shown that GA-rich motifs mediate targeting of *RAB13* mRNA targeting to the actin rich cellular protrusions^37, 38^. Similarly, we have shown that GA-rich localization elements (LEs) in mouse *Net1* mRNA can mediate *Net1* mRNA targeting to neurites and the basal pole of epithelial cells^39^. Our RNAseq analysis of purified MBs showed that there is a significant overlap between mRNAs targeted to the MB, neurite, and basolateral domains of epithelial cells. Furthermore, *RAB13* and *NET1* mRNAs are also both enriched in the MB, suggesting that their 3′ UTR GA-rich motifs may also mediate mRNA targeting to the MBs. Consistent with this hypothesis, we show that the LE present in the 3′ UTR of mouse *Net1* mRNA is necessary and sufficient to target mRNA to the MB. We recently found that these GA-rich motifs likely target RNAs to neurites and the basal pole of epithelial cells by promoting their transport to the plus ends of microtubules in a kinesin 1-dependent manner^40^. This type of transport would be perfect for targeting mRNAs to the MB as plus-ends of microtubules are oriented toward the MB, and kinesin 1 has already been implicated in transporting endosomes to the MB during cell division^41^. All these findings hint at a common mechanism that regulates the subcellular targeting of mRNAs across a variety of cell types and subcellular destinations.

Taken together, this is the first study to demonstrate that a specific subset of mRNAs are targeted and enriched at the MB during cell division and play a role in MB function during abscission (Figure 8). At least some of these mRNAs, such as *CHMP4B* mRNA, appear to be translated at the MB, and this translation plays a key role in accumulation of ESCRT-III components at the MB, thus regulating the abscission step of cytokinesis. While working out the details governing mRNA targeting and translation at the MB will require future studies, this work lays a foundation for the identification of a novel mechanism regulating abscission. Finally, this study will not only advance our understanding of the machinery governing ESCRT-III translocation and accumulation at the MB, but it could also provide novel insight into the mechanisms of RNA targeting to other subcellular compartments and other cell types, as well as providing a framework for studying the role of mRNAs in the post-mitotic MB function.

## Materials and Methods

### Cell culture

All HeLa cell lines were kept in a 37 °C humidified incubator at 5% CO2, routinely tested for mycoplasma, and were maintained in DMEM with 10% FBS and 1% penicillin/streptomycin.

### Cell synchronization

HeLa cells were grown to ∼50% confluency and incubated with 5 mM Thymidine for 16 hours. Cells were then washed and incubated with regular serum-containing media, filled by incubation with 0.1 ug/ml of nocodazole for 16 hours. Mitotic cells were then isolated by gently tapping tissue culture plates to dislodge loosely attached cells arrested in prometaphase. Cells were then washed, seeded on collagen-coated coverslips and incubated in serum containing media for 90 minutes in the presence or absence of puromycin (for last 30 minutes).

### Reagents and antibodies

iTaq Universal SYBR Green Supermix was purchased from Bio-Rad. TRIzol, Puromycin, Doxycycline, Lipofectamine 2000, and Lipofectamine RNAiMax was purchased from Thermo Fisher Scientific. Stellaris Hybridization, Wash A, and Wash B Buffers were purchased from Biosearch Technologies. Hoechst stain was purchased from AnaSpec. Antibodies against the following proteins were used: RPL3 (Bethyl Laboratories #A305-007A,1:100 for immunofluorescence), Acetylated-Tubulin (Cell Signaling #D20G3, 1:500 for immunofluorescence), Acetylated-Tubulin (Millipore Sigma #T7451, 1:500 for immunofluorescence), CHMP4B (Proteintech #16639-I-AP, 1:300 for immunofluorescence), CEP55 (Abnova #H00055165-A01, 1:500 for immunofluorescence), RFP (Chromotek #6g6-20, 1:2000 for immunoblotting), Beta-Actin (LI-COR #926-42210, 1:2000 for immunoblotting), donkey anti-mouse-IgG conjugated to Alexa Fluor 488 (Jackson #715-545-150, 1:100 dilution for immunofluorescence), donkey anti-mouse-IgG conjugated to Alexa Fluor 594 (Jackson #715-585-1500, 1:100 dilution for immunofluorescence), donkey anti-rabbit-IgG Alexa Fluor 488 (Jackson #711-585-152, 1:100 dilution for immunofluorescence), and donkey anti-rabbit-IgG conjugated to Alexa Fluor 594 (Jackson #711-585-152, 1:100 dilution for immunofluorescence), donkey anti-mouse IRDye 680CW (LI-COR #926-32212, 1:5000 for immunoblotting), and donkey anti-rabbit IRDye 800CW (LI-COR #926-68072, 1:5000 for immunoblotting).

### RNA isolation

RNA was extracted using Trizol (Thermo Fisher #15596026) per the manufacturer’s instructions. The RNA was reverse transcribed using SuperScript IV (Thermo Fisher, #18091050) per the manufacturer’s instructions.

### RT-qPCR

Quantitative PCR was performed using iTaq Universal SYBR Green Supermix (Bio-Rad, #1725121). Primer sequences can be found in Supplemental Table 2. To quantify the RT-qPCR, each cDNA sample was normalized to GAPDH. If the experiment was looking at enrichment in MBsomes, the data is represented as a ratio of the normalized MB/normalized total cell. Taqman qPCR reactions were performed using Taqman Fast Advanced Master Mix (Life Technologies) with labeled probe sets (Supplemental Table 3). The quality of each Midbody isolation was quantified by the enrichment of NET1 in MBsomes using the ratio of NET1 / TSG101 in MBsomes compared to NET1/TSG101 in whole cells. Reporter enrichment in MBsomes was quantified using Renilla luciferase as internal control using the ratio of Firefly luciferase / Renilla luciferase RNA in MBsomes compared to the ratio of Firefly luciferase / Renilla luciferase RNA in whole cells.

### MB purification and RNAseq

Midbodies were isolated as previously described^26^. Briefly, media from the various HeLa cells was collected and subjected to a series of centrifugation spins (300 × g, 10,000 × g). Sucrose gradient fractionation was performed at 3000×g, and the interphase between 40% glycerol and 2 M sucrose was collected, spun at 10,000 × g to pellet the midbodies. RNA was extracted using TRIzol (Thermo Fisher #15596026) per the manufacturer’s instructions. For control and MB group 1, the library prep and RNAseq was performed by the University of Colorado Anschutz Medical Campus Genomics facility. The Genomics facility generated a poly A enriched library from the total RNA extraction and performed RNAseq on an Illumina HiSeq2000 platform, utilizing 1 lane, single read 125 cycles. For control group and MB group 2 and 3, the library prep and RNAseq was performed by Novogene. Briefly, Novogene generated a poly A enriched library from the total RNA extraction and performed RNAseq on a NovaSeq PE150 platform with 6 G raw data per sample.

### RNAseq data processing

Transcript abundances were calculated using Salmon v1.9.0^42^ using the flags –seqBias and – gcBias. Transcript abundances were collapsed to gene level using txImport [cite: https://f1000research.com/articles/4-1521/v1]. To determine enrichment of genes in the midbody, DEseq2 was used to calculate the log2 ratio of midbody/whole cell counts and identify significantly enriched RNAs^49^. Genes with a basemean (average of normalized count values) less than 250 were excluded. Plots were generated in Rstudio using R version 4.1.1.

### siRNA transfections

Custom CHMP4B siRNA oligonucleotides (Dharmacon, AUCGAUAAAGUUGAUGAGUUAUU) and custom CEP55 siRNA oligonucleotides (Millipore Sigma, AGGCAUGUACUUUAGACUU) were transfected into HeLa cells using Liptofectamine RNAiMAX with 40nM oligonucleotide. The efficiency of the knock-down was measured 72 h post transfection by RT-qPCR (previously described above).

### Immunofluorescence and quantification

HeLa cells were treated as indicated in the text and then fixed in 1% paraformaldehyde in PBS for 10 min at room temperature. Cells were rinsed three times in PBS. The cells were then incubated with primary antibody in PBS containing 0.5% BSA and 0.2% saponin for 1 h at room temperature, washed three times in PBS and then incubated with the appropriate fluorochrome-conjugated secondary antibodies diluted in PBS containing 0.5% BSA and 0.2% saponin for 30 min. Cells were washed three times in PBS and mounted in Fluoromount (Southern Biotech).

Fixed cells were imaged with an inverted Axiovert 200M microscope (Zeiss) with a × 63 oil immersion lens and QE charge-coupled device camera (Sensicam). Z-stack images were taken at a step size of 500–1000 nm. Image processing and quantification was performed using 3i Slidebook 6 software (Intelligent Imaging Innovations). Briefly, masks were made to calculate the total intensity of the cell body or the intercellular bridge as well as the total area for each mask. The numbers were then used to calculate the intensity/micron^2^. Unless otherwise stated, all images are represented as a maximal projection of the z-stack.

### Time-lapse imaging

HeLa cells stably expressing GFP-MKLP1 were plated on 6cm glass-bottom dishes and imaged at 37^0^C for 120 minutes with time lapse of 15 minutes (to minimize phototoxicity). To ensure that cells are in mid-late telophase cells were selected based on following criteria: cells were flattened with round nucleus (kidney-shaped nucleus is a feature of cells in early telophase) containing identifiable MB (labeled by GFP-MKLP1). Where indicated, 25μM puromycin was added at the beginning of time-lapse analysis.

### Proximal Ligation Assay

HeLa cells stably expressing GFP-MKLP1 were transfected with either HA-CHMP4B or HA-CHMP4B-3′UTR constructs. Non-transfected HeLa cells were used as one of the negative controls. Cells were then treated with puromycin for 10 minutes (not treated cells were used as another negative control), followed by fixation with 4% paraformaldehyde and permeabilization with 0.1% Triton X-100. Cells were incubated with anti-HA and anti-puromycin antibodies, followed by Duallink proximal ligation assay (PLA) as described in manufacturers protocol (Sigma). For imaging, cells in mid-to-late telophase were randomly picked using following criteria: (1) cells had a clearly identifiable MB as determined by GFP-MKLP1 signal; (2) cells had fully reformed round nuclei; (3) cell bodies were flattened; (4) daughter cells were connected by extended intercellular bridge of at least 2 μm in length.

### smiFISH probe design

The smiFISH protocol was adopted from Tsanov *et al*^27^. Briefly, probes were designed by inputting the gene sequences (CHMP4B, NET1, TSG101) into the Oligostan software (Supplemental Table 4). smiFISH probes then had a Y flap (TTACACTCGGACCTCGTCGACATGCATT) added in order to have a location where we can hybridize the fluorescent molecules to the probe. The sequences can be found in Supplemental Table 4. The fluorescent probe was designed by taking the reverse complement of the Y flap (AATGCATGTCGACGAGGTCCGAGTGTAA) and adding Alexa488, Alexa594, or Cy3 to both ends.

### smFISH probe design

smFISH probes for Firefly luciferase were obtained from Biosearch Technologies. The probes were labeled with Quasar 570 dye. The sequences of these probes are proprietary.

### smiFISH/smFISH visualization

HeLa cells were plated on collagen coated coverslips and allowed to grow to 70-75% confluency. The media was aspirated, and cells were washed once with 1x PBS. For Halo-tag visualization, HaloTag Oregon Green Ligand was added 4-6 hours prior to fixing cells. Cells were fixed in 1% PFA for 10 min at room temperature, then washed twice with 1x PBS. Cells were permeabilized with 0.1% TX-100 for 5 min at room temperature. For Halo-tag visualization, cells were not permeabilized to preserve Halo-tag fluorescence. The cells were washed with Wash Buffer A (Stellaris) at room temperature for 5 min. In the meantime, the CHMP4B probes and Y flap was hybridized using the protocol from Tsanov *et al*^27^. For smFISH, this hybridization was not necessary. 2μl of the probe/flap hybridization product (0.833µM) or smFISH probe was added to 100μl of smFISH hybridization buffer (Biosearch Technologies). A hybridization chamber was prepared using an empty 15cm cell culture plate, wrapped in tinfoil with wet paper towels surrounding the rim on the inside and parafilm covering the bottom of the plate. 100μl of the probe-containing hybridization solution was added to the parafilm. The coverslip was then placed on top of this droplet of hybridization buffer with the cell side down. The hybridization chamber with the coverslips was incubated at 37°C overnight. On the following day, the glass coverslips were transferred to a fresh 6-well plate with the cell side up and incubated with Wash Buffer A (Biosearch Technologies) and Hoechst stain (1:2000, AnaSpec) at room temperature in the dark for 30 min. Hoechst stain was washed with Wash Buffer B (Biosearch Technologies) at room temperature in the dark for 5 min. Coverslips were then mounted onto slides with Fluoromount G (Southern Biotech) and sealed with clear nail polish. Slides were imaged using an inverted Axiovert 200M microscope (Zeiss) with a 63X oil immersion lens and QE charge-coupled device camera (Sensicam).

### smFISH particle counting

ImageJ was used to count the total number of smiFISH particles. First, the channels were separated, and an outline of the cell was drawn using the freehand tool in the smFISH channel. Next, a threshold was set using a control cell. The threshold set with the control cell was used for all cells after that. Finally, the particles were analyzed to get the total number of particles in each cell.

### FISH-quant

FISH-quant was used to quantify midbody or intercellular bridge enrichment of smFISH spots as previously described^50^. Briefly, outlines were drawn in the FITC channel visualizing Halo-MKLP1. Four outlines were drawn per dividing cell: 2 cells on each side of the bridge, the intercellular bridge, and the midbody marked by Halo-MKLP1. Prior to quantification, identified smFISH spots were thresholded for intensity, sphericity, amplitude and position. Reporter enrichment was quantified by the total number of spots in the intercellular bridge or midbody of cells over total number of spots in the whole cell normalized to the control.

### Puromycin treatment

HeLa cells were plated on collagen-coated coverslips and allowed to grow to 70-75% confluency. At this time, cells were exposed to 25μM puromycin and incubated at 37°C for 1 h. Cells were then fixed, stained, and imaged as previously described.

### Anti-Puromycin labeling

HeLa cells were plated on collagen-coated coverslips and allowed to grow to 70-75% confluency. At this time, the cells were either treated with DMSO (volume equal to that of the other treatments) for 40 min at 37°C, 50μM cycloheximide for 40 min plus the last 10 min with 25μM puromycin at 37°C, or only 25μM puromycin for 10min at 37°C. Cells were then fixed, stained, and imaged as previously described.

### Puromycin Wash-Out Assay

HeLa cells were plated on collagen-coated coverslips and allowed to grow to 70-75% confluency. At this time, the cells were either untreated, treated with 25μM puromycin at 37°C for 1 h, treated with 25μM puromycin at 37°C for 1 h, washed with 1x PBS and allowed to recover for 5 min in complete media at 37°C, or treated with 25μM puromycin at 37°C for 1 h, washed with 1x PBS and allowed to recover for 10 min in complete media at 37°C. Cells were then fixed, stained, and imaged as previously described.

### Generation of doxycycline-inducible constructs

Due to the GC rich nature, we were unable to PCR the full-length 3’-UTR. Therefore, we used Twist Bioscience to generate a modified sequence that we could use for a template (TGGGGTCCAGCGCTGGCTGGGCCCAGACAGACTGTGGTGGCCTGCGCAGCGAGCAGGCGTGTGCGTGTGTGGGGCAGGCAGGATGTGGTGCAGGCAGGTTCCATCGCTTTCGACTCTCACTCCAAAGCAGTAGGGCCGCGTTGCTGCTCACTCTCTGCATAGCATGGTCTGGAAGTGTGCTGTTTATAATGTTGAATTTCTGTAAAATAAACTGTATTTGCAAATCCAACATTGAGCTTCTGGACTACGCTGACTCCACTGCTGAATCCTCAATGGAAAGGGTCGACTGGTTGCAGTTGAAATGACCTGAAATGTAGCCTCTGTCCTTGTAAGTCAGTTGACTTGCCGCACATCTCTTTGTGTACTTGTACGGTACTGGCAGAAAAGTCATTTTTCAAAAGCCATAGGCTTTTCCTTGCCCTTAGCTGTAATAATGCATCTGATTTTGATTTCCTCCAGAGCTGTGTTTCTGTCCATCACCTGTGTATTGGCCCTGTGTTTACCACTCTGGCCCACTCCTCACCCCCTTGCTCCCCTGGTCTTCTGGAGTTTGTGACATTGATTTGAAATGGATGGTGTTCTCTTGAGAGCAAGTGAGATTGTTAGAATTAAGTTCCAACTATACAGTTTTCTAACATAGCTATAAGGTCCTTGTTGCTGTTTGTGATAACTGATAGATAACTCATTGGAAACGTGCATACATTTATATTCAGATGAAATTATGGTTTGCACTGTCTATTAAATATCTCGATTAATTTTCATA). This sequence maintains the GA rich motifs thought to be important for translocation.

To insert the various doxycycline-inducible constructs into the backbone (pRD-RIPE), we used the NEBuilder Assembly tool and followed the programs direction to generate forward and reverse primers for generating PCR products for the insert and the backbone (listed in the Supplemental Table 5). Q5 High-Fidelity 2x Master Mix (New England Biolabs) was used to perform the PCR according to the manufacturer’s protocol. NEBuilder HiFi DNA Assembly Master Mix was used to assemble the inserts and backbones according to the manufacturer’s protocol and transformed into XL10-Gold competent cells (Agilent). Bacteria containing the constructs were then grown overnight at 37°C and the DNA was extracted using the Zyppy Plasmid Miniprep Kit (Zymo Research) according to the manufacturer’s protocol.

For the RFP-3′UTR, RFP-CHMP4B-3′UTR were generated as described above and then using the NEBuilder system, the CHMP4B coding region was removed by PCR and re-assembled to get RFP-3′UTR construct. For the RFP-CHMP4B No Start, the RFP-CHMP4B was generated as described above and then using the NEBuilder system, the start codon was removed by PCR and re-assembled to get the RFP-CHMP4B No Start construct.

### Transfection of doxycycline-inducible constructs

HeLa acceptor lines containing a HeLa LoxP site were plated in a 6-well plate and allowed to grow to 60% confluency. Cells were then co-transfected with a pRD-RIPE construct containing a gene of interest, along with a Cre-encoding plasmid (pBT140) at a 0.5%-10% wt/wt ratio. To transfect 1 well in a 6-well plate, a total of 2.1μg of DNA was mixed with 5.625μL of Lipofectamine 2000 (Thermo Fisher Scientific) and 246μL of Opti-MEM (Invitrogen) following the manufacturer’s protocol. Cells were incubated with the transfection mixture overnight, the medium was changed, and the incubation continued for another 24 h before adding puromycin. We used a two-step selection protocol, beginning with half of the maximal puromycin concentration (0.5μg/mL) for the first 48 h of selection, followed by the maximal puromycin concentration (1μg/mL) for several days until the puromycin-sensitive cells were eliminated. The cultures were incubated until the appearance of visible puromycin-resistant colonies, which were then pooled together and expanded.

### Induction of doxycycline-inducible cell lines

Cells containing a doxycycline-inducible gene of interest were subject to 2μg/mL of doxycycline and allowed to grow for 48 h unless otherwise stated. After 48 h, the cells were either fixed on collagen-coated coverslips for immunofluorescence, lysed for western blot analysis, or the media was collected for MBsome isolation.

### Cell lysis and western blot analysis

Cells were lysed on ice in a buffer containing 20 mM Hepes, pH 7.4, 150 mM NaCl, 1% Triton X-100, and 1 mM PMSF. After 30 min, lysates were clarified at 15,000 g in a prechilled microcentrifuge. Supernatants were collected and analyzed via Bradford assay (5000006; Bio-Rad Protein Assay). 50-µg lysate samples were prepared (unless otherwise stated) in 4× SDS loading dye, boiled for 5 min at 95°C, and separated via SDS-PAGE. Gels were transferred onto 0.45-µm polyvinylidene difluoride membrane (IPFL00010), followed by blocking for 30 min in Intercept Blocking Buffer diluted in TBST 1:3 (927-60001). Primary antibodies (made in diluted Intercept Blocking Buffer) were incubated overnight at 4°C. The next day, blots were washed in TBST followed by incubation with IRDye fluorescent secondary antibody (diluted Intercept Blocking Buffer) for 30 min at room temperature. Blots were washed once again with TBST before final imaging on a Li-Cor Odyssey CLx.

### Statistical analysis

All statistical analyses were performed using GraphPad Prism Software (GraphPad, San Diego, CA). A Student’s t-test was used to determine significance unless otherwise noted. Error bars represent standard deviation unless otherwise noted. For all immunofluorescence experiments, at least 20 randomly chosen image fields from at least 3 separate experiments were used for data collection. For quantitative immunofluorescence analysis, the same exposure was used for all images in that experiment and was quantified using Intelligent Imaging Innovations software (Denver, CO, USA). For western blot experiments, 3 separate experimental examples were used for data collection. For quantitative western blot analysis, Image Studio V5.2 software (LI-COR, Lincoln, NE) was used to collect intensities.

## Acknowledgements

We would like to thank the University of Colorado Anschutz Medical Campus Genomics Facility and Novogene for performing the RNAseq and the RNA Biosciences Initiative (RBI) for help in analyzing the RNAseq data and Migle Prekeryte for critically reading and editing the manuscript. This work was supported by NIGMS grant R01-GM143774 to RP, R35-GM133385 to JMT, and RBI Pilot Grant to RP and JMT. Additionally, some of the work was also supported by Diversity Supplement GM143774-02S1 to KV.

---

**Supplemental Figure 1.**

(A) Comparison of MB-Associated mRNA with RNA contents of HeLa cells in G2/M and G1 cell cycle phases. (B) Comparison of MB-Associated mRNA with RNA contents of HeLa cells in S and G1 cell cycle phases. (C) Comparison of MB-Associated mRNA with spindle-associated RNA.

**Supplemental Figure 2.**

(A-B) GFP-MKLP1 expressing HeLa cells were either mock treated or treated with CHMP4B (A) or CEP55 (B) siRNAs. Cells were then incubated for 72 h, followed by mRNA isolation and RT-qPCR analysis using CHMP4B, CEP55, and GAPDH (control) specific primers. Data shown are the CHMP4B or CEP55 mRNA levels normalized against GAPDH mRNA.

(C-D) GFP-MKLP1 expressing HeLa cells were either mock treated (top two panels) or treated with puromycin for 1 h (bottom two panels) and subject to staining with smFISH probes against CHMP4B mRNA. CHMP4B mRNA particles are marked by the arrows in the inset and the black dashed square represents the region of the image used for the inset (C). The number of cells with CHMP4B smFISH signal was then counted. The data shown in panel D represents the means and standard deviations derived from three independent experiments.

(E) GFP-MKLP1 expressing HeLa cells were either mock treated or treated with CEP55 siRNA for 72 hours. Cells were then fixed and subjected to staining with smFISH probes against CHMP4B mRNA. The number of cells with CHMP4B smFISH signal in the MB were then counted and expressed as means and standard deviations derived from three independent experiments.

**Supplemental Figure 3. The effect of puromycin treatment on CHMP4B targeting to the MB in synchronized cells**

(A) Synchronization and puromycin treatment schematics

(B-C) Synchronized HeLa cells were incubated for 90 minutes in the presence or absence of puromycin (during last 30 minutes of incubation). Cells were then foxed and stained with anti-CHMP4B antibodies (red). Asterisk marks the midbody. Panel C shows quantification of CHMP4B localization in the midbody. Data shown are the means and standard deviations. Dots represents localization in individual cells.

**Supplemental Figure 4.**

(A-B) HeLa cells were fixed and subjected to immunostaining using anti-acetylated-tubulin. The panel A represents a cell in telophase. Panel B represents a cell that had just undergone abscission (see arrow). The lines used for the line intensity graphs are marked in yellow, and the white dashed square represents the region of the image used for the inset.

(C-D) Line intensity graphs representing the intensity of acetylated-tubulin in telophase and abscission cells. The abscission site is marked by an arrow (D).

(E-F) Telophase HeLa cells stably expressing GFP-MKLP1 were analyzed by time-lapse microscopy. Cells were imaged for 120 minutes with 15 minute time-lapse.

**Supplemental Figure 5. Characterization of HeLa LoxP cells expressing doxycycline-inducible RFP-tagged constructs.**

(A) HeLa LoxP cells expressing various RFP-tagged dox-inducible constructs cells were untreated (-Dox) or treated with 2μg/mL doxycycline for 48 h (+Dox). RFP was visualized by fluorescent microscopy to test for dox-dependent expression. The white dashed square represents the region of the image used for the inset.

(B-D) HeLa LoxP cells expressing various RFP-tagged dox-inducible constructs cells were untreated (-Dox) or treated with 2μg/mL doxycycline for 48 h (+Dox). The construct expression levels were then determined by RT-qPCR using CHMP4B specific primers.

(E) HeLa LoxP cells expressing various RFP-tagged dox-inducible constructs cells were treated with 2μg/mL doxycycline for 48 h. The levels of expression among various constructs were the n compared using RT-qPCR with RFP specific primers.

(F-G) HeLa LoxP cells expressing various RFP-tagged dox-inducible constructs cells were untreated (-Dox) or treated with 2μg/mL doxycycline for 48 h (+Dox). The levels of RFP-tagged protein expressions were visualized via western blot using anti-RFP and anti-Actin (loading control) antibodies. Panel G shows Western blot quantification from 3 individual experiments. Statistical analysis is represented with a p-value.

**Supplemental Figure 6.**

(A) smiFISH for endogenous *NET1* mRNA localization (red) in human epithelial cells. As a nonlocalized control, transcripts encoding an exogenous Firefly luciferase are also visualized.

(B) Quantification of *NET1* RNA localization along the apicobasal axis of epithelial cells.

(C) Quantification of MB-localized *Net1* UTR-containing reporter transcripts. In all samples, reporter transcript counts in midbodies and whole cells were quantified and the ratio of counts between the two locations is reported. These ratios were normalized by setting the value for the control reporter transcript lacking 3′ UTR additions to one. P values were calculated using a t-test.

**Supplemental Table 1. List of mRNAs identified in RNAseq analyses of purified MBs**

**Supplemental Table 2. SYBR RT-qPCR primer sequences**

**Supplemental Table 3. Taqman RT-qPCR primer sequences**

**Supplemental Table 4. smiFISH probe sequences**

**Supplemental Table 5. Primer sequences used for cloning**

