## Supplementary figures and images for "The Novel Role of Midbody-Associated mRNAs in Regulating Abscission"

### Supplemental Figure 1

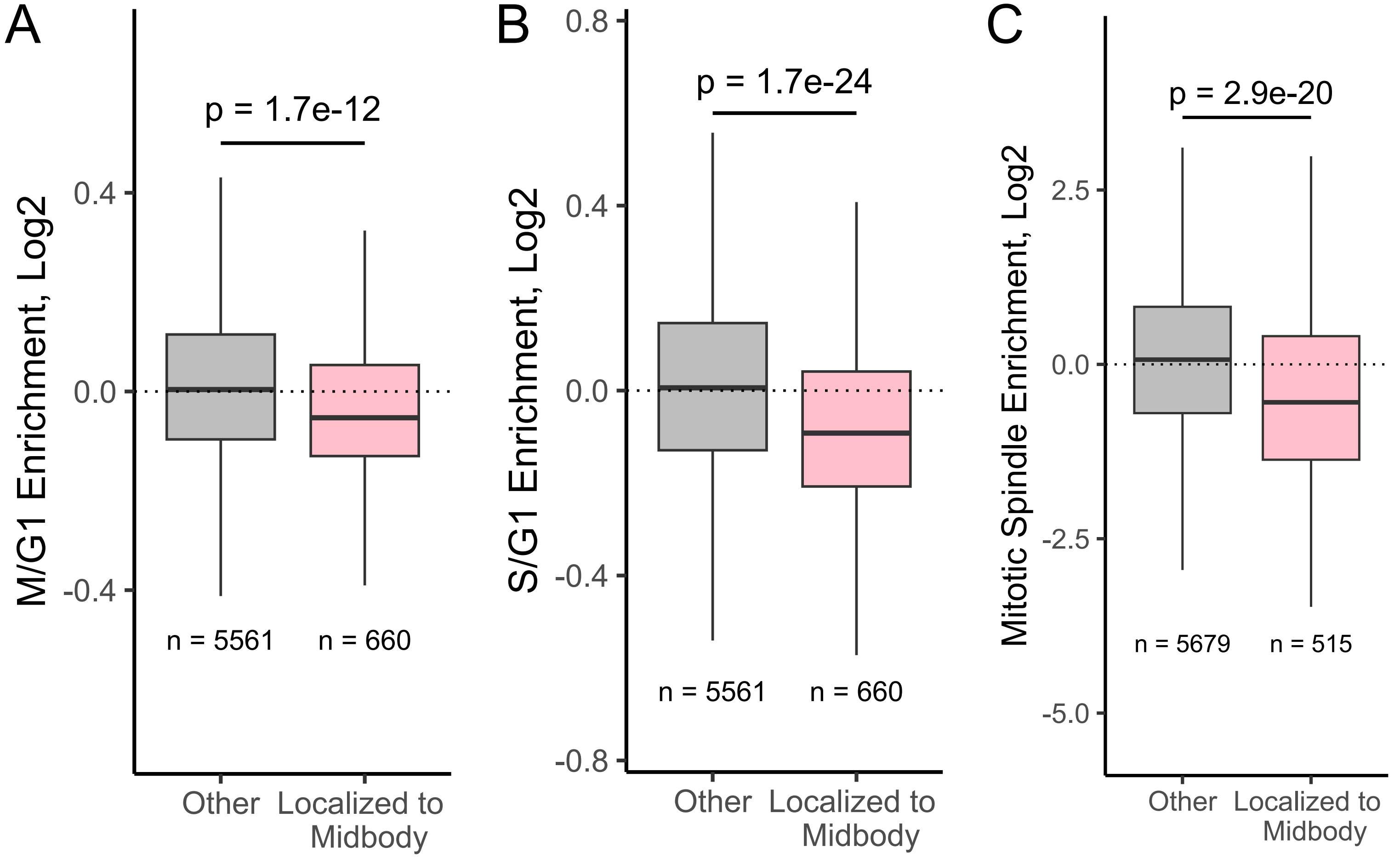

### Supplemental Figure 2

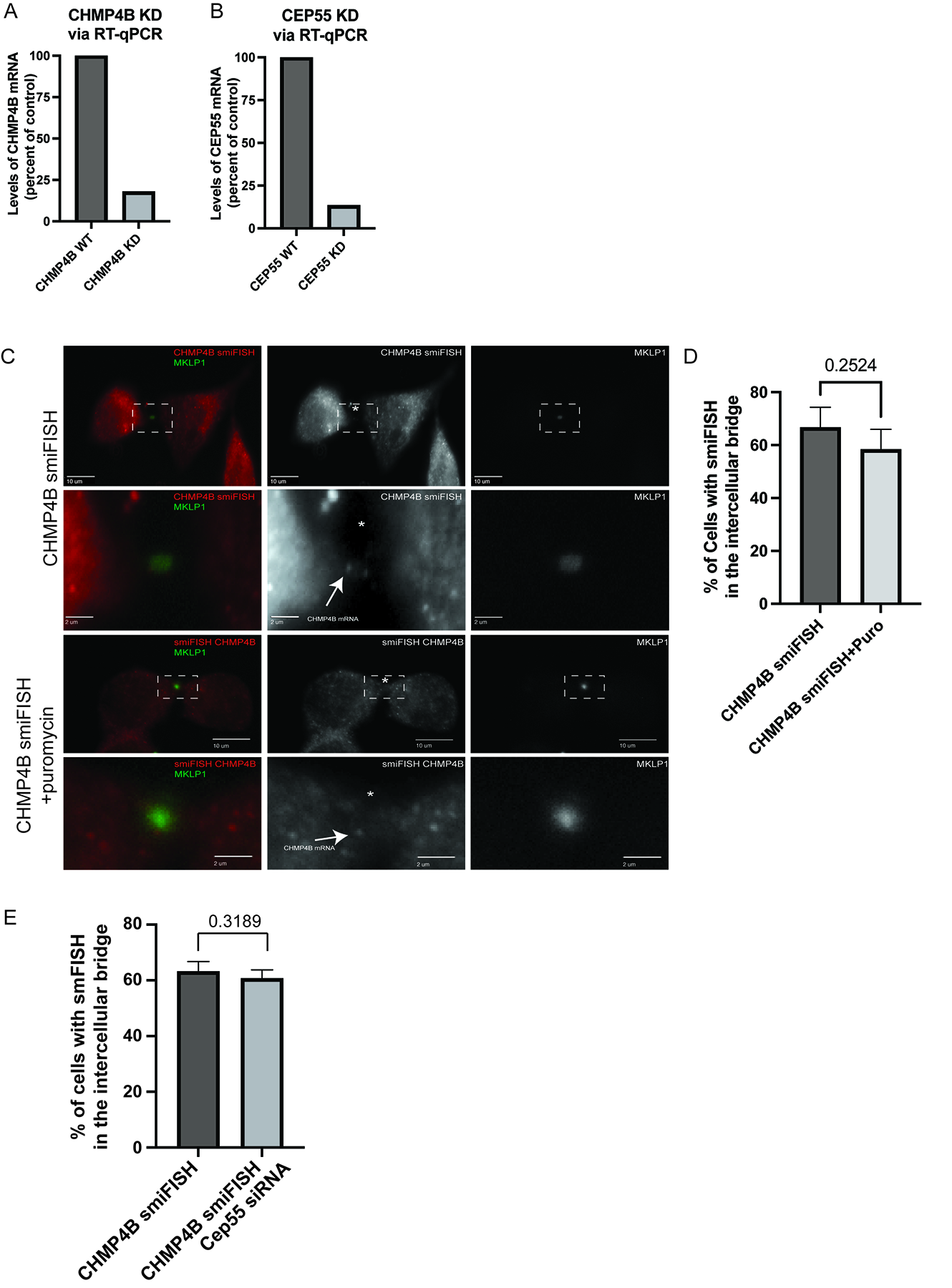

### Supplemental Figure 3

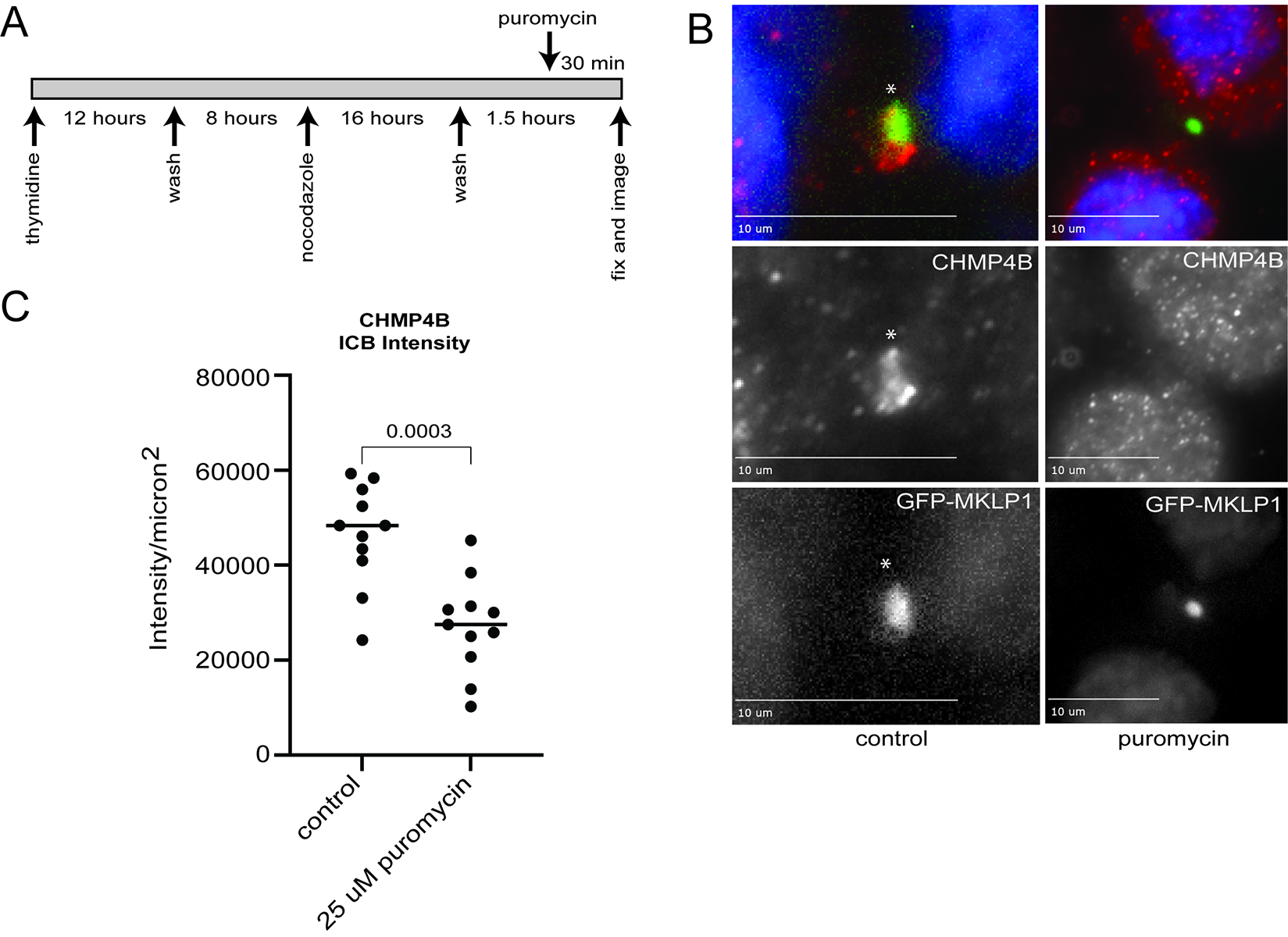

### Supplemental Figure 4

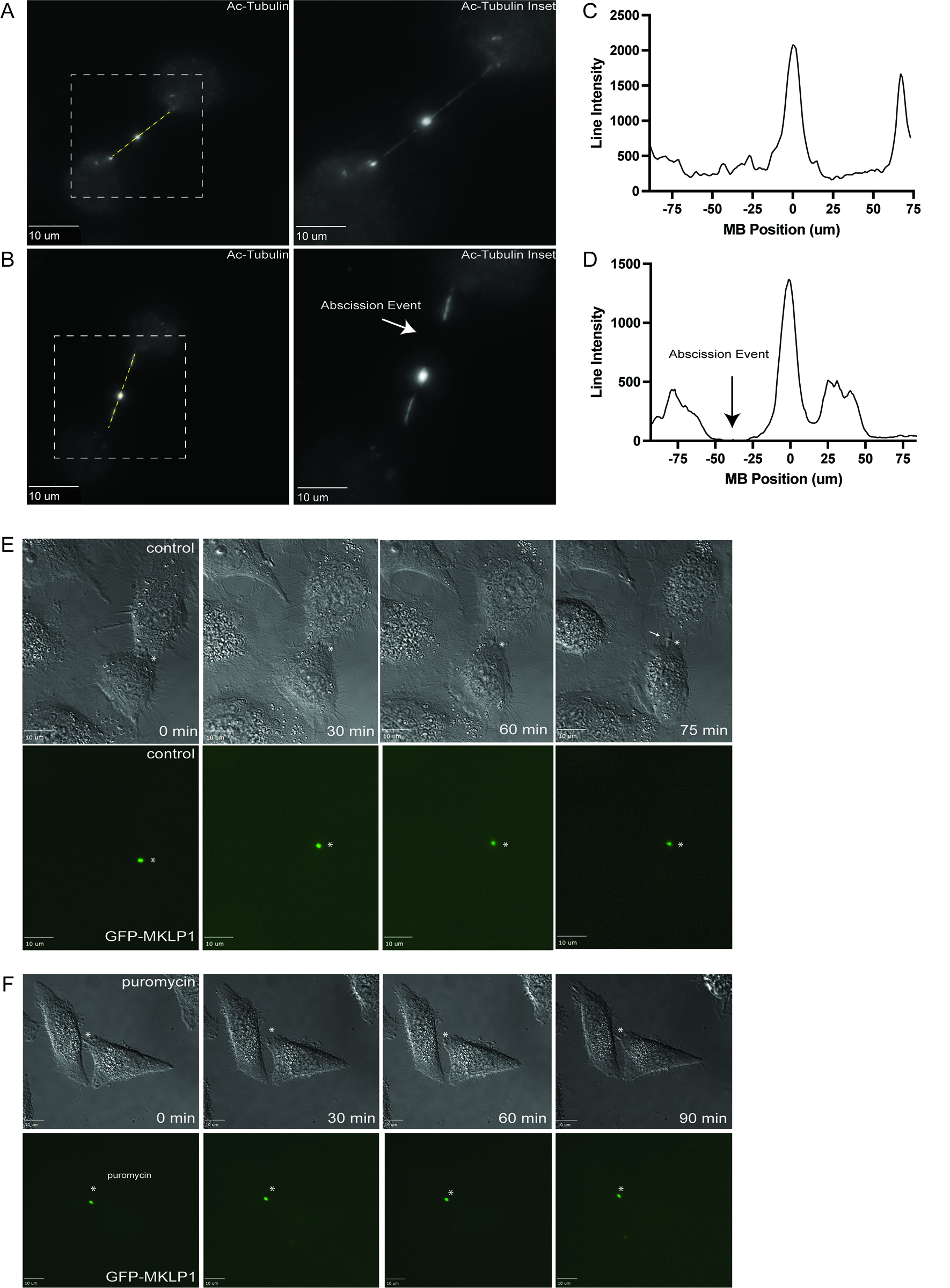

### Supplemental Figure 6

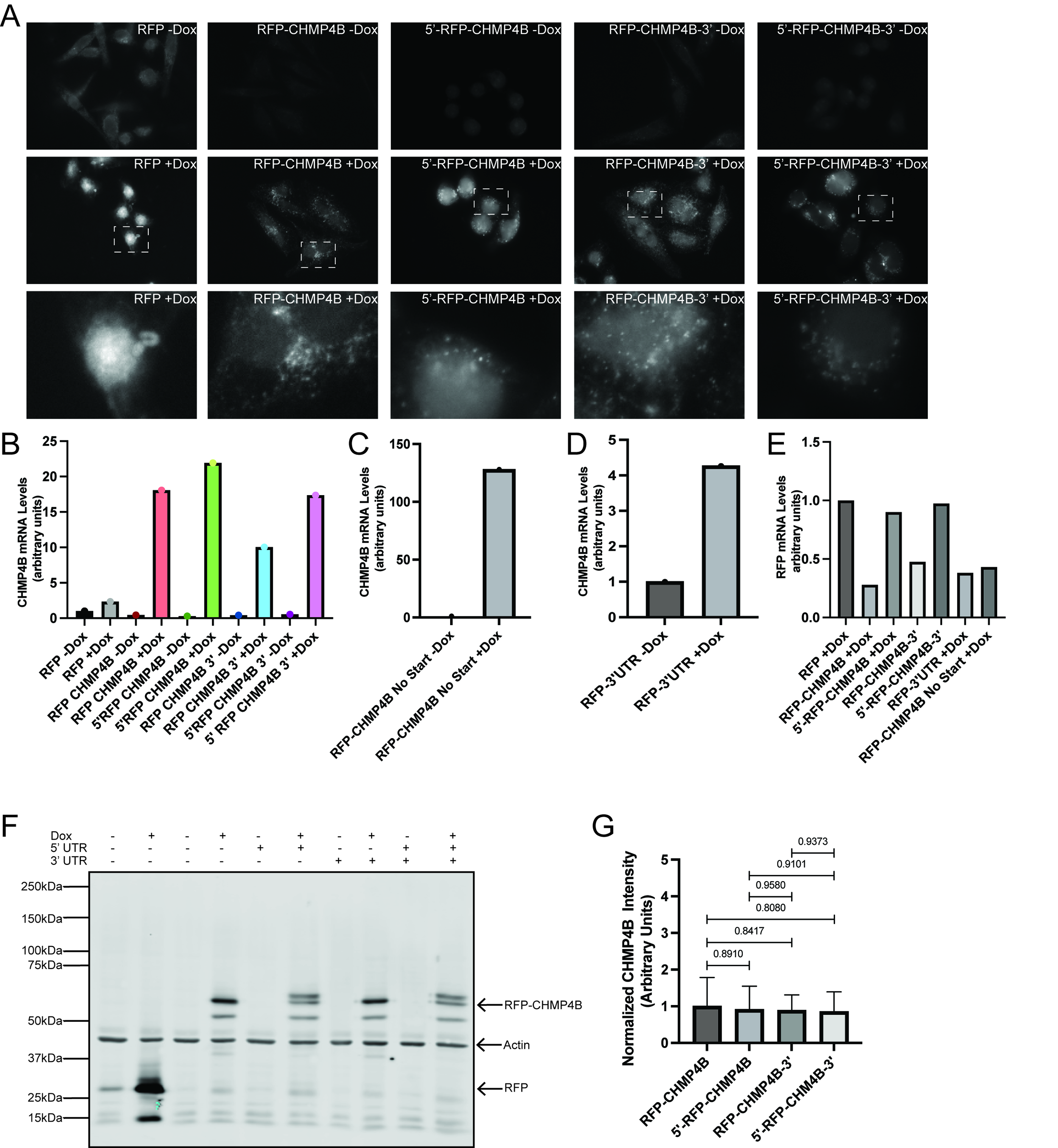

### Supplemental Figure 7

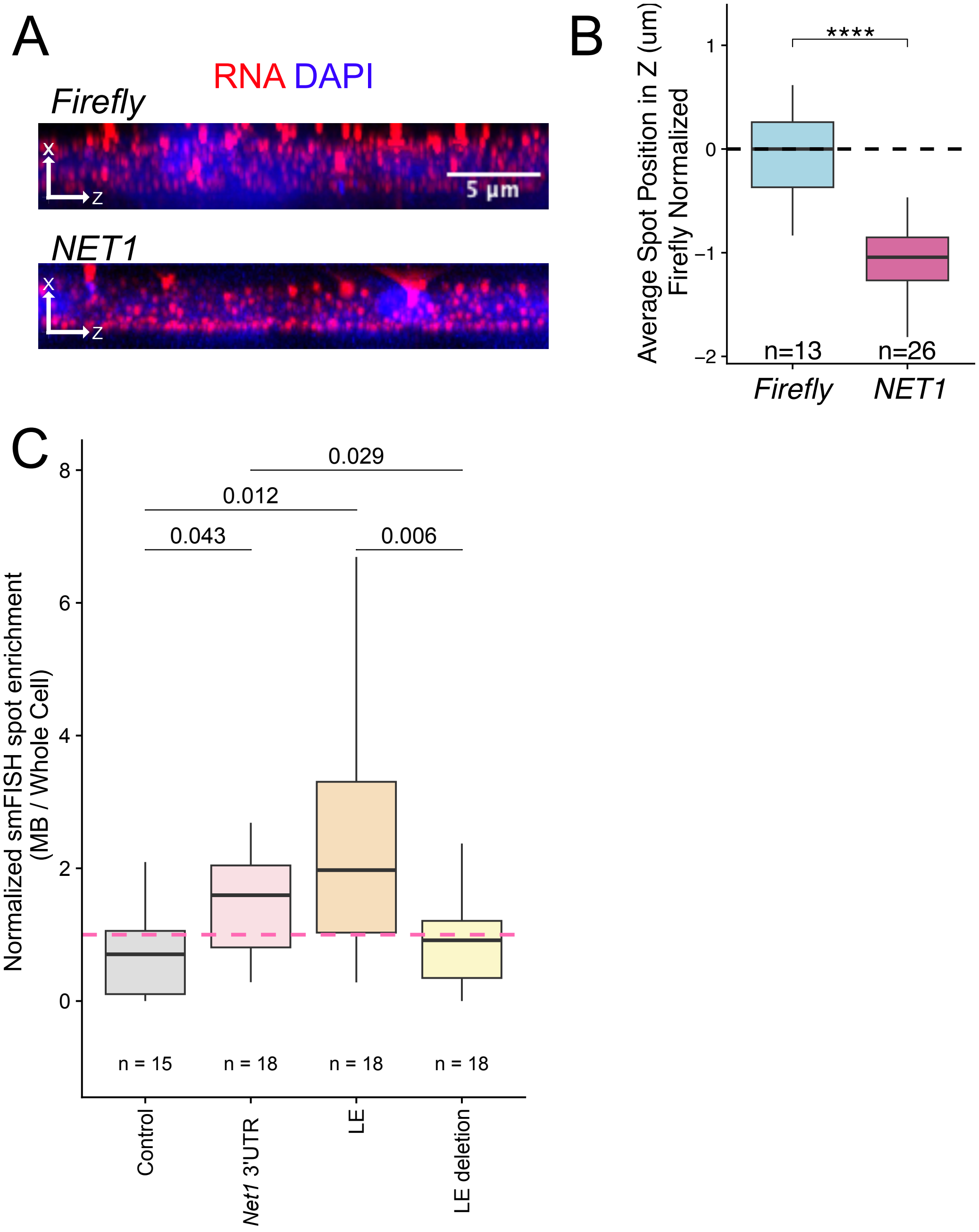
